# Single-cell and Spatial Transcriptomics Identified Fatty Acid-binding Proteins Controlling Endothelial Glycolytic and Arterial Programming in Pulmonary Hypertension

**DOI:** 10.1101/2024.02.11.579846

**Authors:** Bin Liu, Dan Yi, Shuai Li, Karina Ramirez, Xiaomei Xia, Yanhong Cao, Hanqiu Zhao, Ankit Tripathi, Shenfeng Qiu, Mrinalini Kala, Ruslan Rafikov, Haiwei Gu, Vinicio de jesus Perez, Sarah-Eve Lemay, Christopher C. Glembotski, Kenneth S Knox, Sebastien Bonnet, Vladimir V. Kalinichenko, You-Yang Zhao, Michael B. Fallon, Olivier Boucherat, Zhiyu Dai

## Abstract

Pulmonary arterial hypertension (PAH) is a devastating disease characterized by obliterative vascular remodeling and persistent increase of vascular resistance, leading to right heart failure and premature death. Understanding the cellular and molecular mechanisms will help develop novel therapeutic approaches for PAH patients. Single-cell RNA sequencing (scRNAseq) analysis found that both FABP4 and FABP5 were highly induced in endothelial cells (ECs) of *Egln1^Tie2Cre^* (CKO) mice, which was also observed in pulmonary arterial ECs (PAECs) from idiopathic PAH (IPAH) patients, and in whole lungs of pulmonary hypertension (PH) rats. Plasma levels of FABP4/5 were upregulated in IPAH patients and directly correlated with severity of hemodynamics and biochemical parameters using plasma proteome analysis. Genetic deletion of both *Fabp4* and 5 in CKO mice (*Egln1^Tie2Cre^/Fabp4-5^-/-^*, TKO) caused a reduction of right ventricular systolic pressure (RVSP) and RV hypertrophy, attenuated pulmonary vascular remodeling and prevented the right heart failure assessed by echocardiography, hemodynamic and histological analysis. Employing bulk RNA-seq and scRNA-seq, and spatial transcriptomic analysis, we showed that *Fabp4/5* deletion also inhibited EC glycolysis and distal arterial programming, reduced ROS and HIF-2α expression in PH lungs. Thus, PH causes aberrant expression of FABP4/5 in pulmonary ECs which leads to enhanced ECs glycolysis and distal arterial programming, contributing to the accumulation of arterial ECs and vascular remodeling and exacerbating the disease.

## Introduction

PAH is a progressive, complex, and devastating disease arising from a variety of pathogenic and genetic causes. Excessive vasoconstriction and abnormal vascular remodeling have been considered as the major factors contributing to the complicated pathogenesis of PAH^1,2^. Owing to the poor understanding of the underlying mechanisms of obliterative vascular remodeling, current therapies result in only modest improvement in morbidity and mortality with 5 years survival rate around 50%^3^. Accumulating evidence shows that PAH is a systemic metabolic disease^4–6^. Previous studies showed that non-esterified free fatty acids and acylcarnitines in the circulation are upregulated in patients with PAH compared to healthy controls, which was linked to RV lipotoxicity^7–10^. Using a combination of high-throughput liquid-and-gas-chromatography-based mass spectrometry analysis on human PAH lung, Zhao et al demonstrated an increase of β-oxidation in fatty acids and upregulation of lipid oxidation in PAH lungs^11^. However, these studies are mainly observational and the role of fatty acid metabolism during PAH development and progression remains elusive^12^.

Blood vessels deliver oxygen and nutrients including lipids. Circulating fatty acids are transported in the forms of triglycerides, which are released by lipoprotein lipase (LPL) anchoring in the lumen side of the endothelium^13^. Mounting evidence demonstrated that fatty acid chaperones mediate transendothelial fatty acid transport^14^. Fatty acid-binding protein (FABP) 4 and 5 are members of intracellular FABPs, which bind long-chain fatty acids and facilitate lipids transport in and between cells. FABP4 is firstly identified in mature adipocytes, while FABP5 is expressed abundantly in the skin epidermal cells^15^. Recent studies showed that FABP4 and FABP5 are expressed in ECs across multiple tissues^16,17^, and circulating FABP4 level is elevated in the PAH patients^18^. However, the role of endothelial FABP4/5 in the pathogenesis of PAH remains undetermined.

Loss of distal capillaries and accumulation of pathogenic pulmonary arterial ECs with proliferative, anti-apoptotic and glycolytic phenotypes are hallmarks of PAH development^19–23^. Our recent studies demonstrated that arterial programming is evident in the distal microvasculature of PAH lungs and PH mice lung due to the general capillary ECs transition to arterial ECs through HIF-2α/Notch4 signaling^24^. In the present study, we found that among fatty acid chaperones, both FABP4 and FABP5 are dramatically induced in mouse PAECs from *Egln1^Tie2Cre^* mice (CKO), a novel mouse model with severe PH and right heart failure^25,26^, and in human PAECs isolated from IPAH patients. Based on scRNAseq and spatial transcriptomics analysis, genetic deletion of *Fabp4/5* dramatically reduced ECs arterialization in the distal lung and glycolytic programming in CKO mice. We aimed to determine how lung endothelial FABP4/5 regulate arterial ECs accumulation in the pathogenesis of PAH.

## Results

### FABP4/5 were upregulated in PAECs from human PAH and rodent PH

We recently performed scRNA-seq analysis to profile pulmonary cells in *Egln1^f/f^* (WT) and *Egln1^Tie2Cre^* (CKO) mice at the age of 3.5 months, at which age the CKO mice develop severe PH^26–28^. Using scRNA-seq analysis^29^, we found that FABP4 was expressed at relatively high levels in ECs and macrophages in native (WT) lungs, whereas FABP5 was expressed in alveolar type 2 cells (AT2), alveolar macrophages (aMac), and ECs in the native lung (**Fig.1a**). *Fabp4* and *Fabp5* expression was significantly higher in ECs but not aMac of CKO mice compared to WT mice (**Fig. 1a**). Immunostaining confirmed that FABP4 and FABP5 were highly expressed in highly remodeled and occlusive vessels of CKO lungs (**Fig. 1b**). Moreover, Western blotting showed a significant increase in FABP4 and FABP5 protein amounts in lungs of PH rodent models including CKO mice, monocrotaline (MCT)^30^ and Sugen5416/hypoxia(SuHx)-induced PH rats^31^ (**Fig. 1c and 1d**). In addition to the animal models, endothelial FABP4/5 expression was also significantly increased in isolated PAECs from patients with IPAH compared to PAECs from healthy donors (**Fig. 1e and 1f**). Our data were consistent with a recent study showing the upregulation of circulating FABP4 in PAH patients and lung *Fabp4* mRNA in PH rodent models^18^. Thus, endothelial FABP4/5 are induced in mouse, rat, and human PAH lungs.

**Figure 1.**
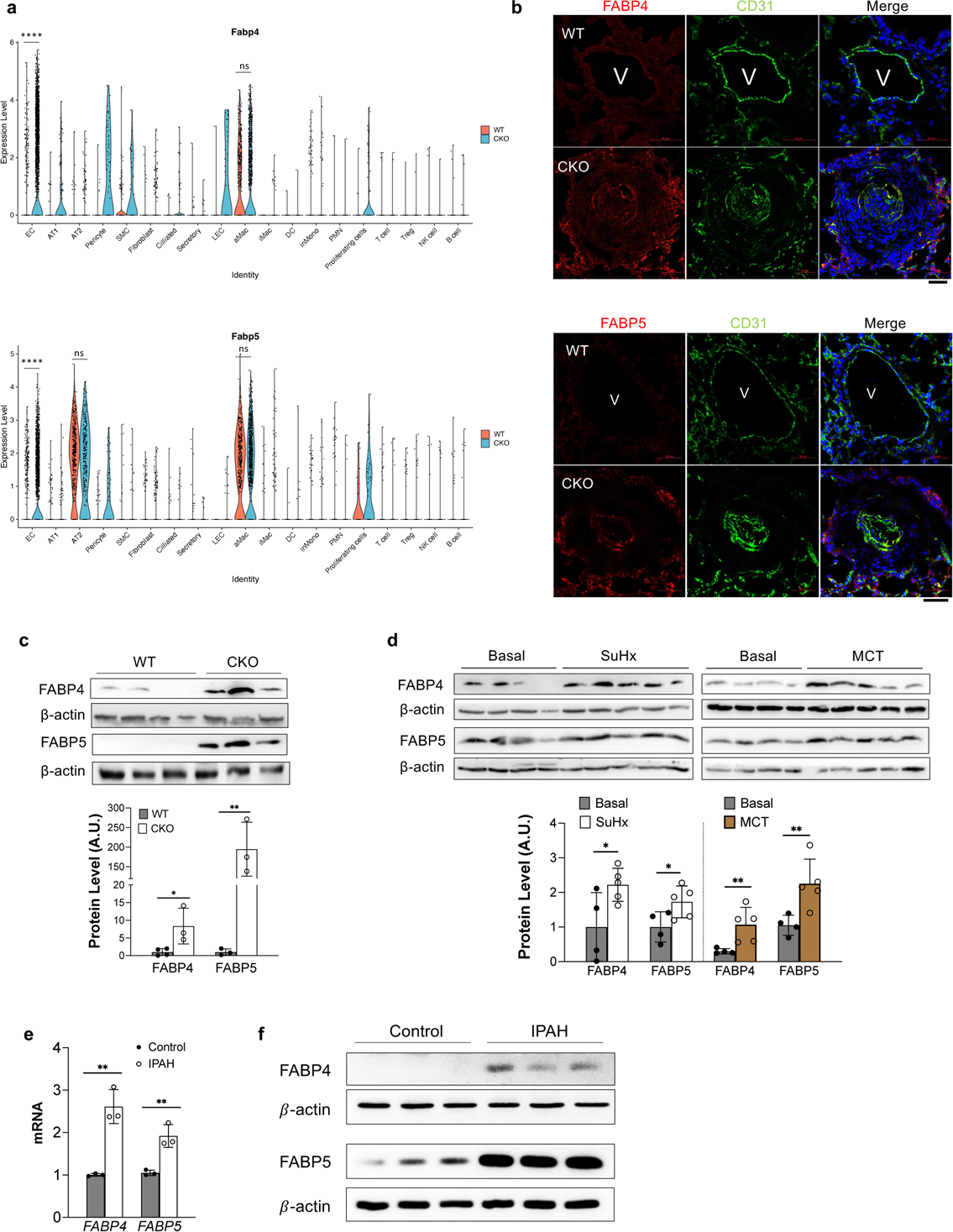
Upregulation of FABP4 and 5 in the PAH patients and PH rodents. **(a)** Single-cell RNA sequencing analysis showed that FABP4 and 5 were upregulated in the lung ECs of *Egln1^Tie2Cre^* (CKO) mice compared to WT mice. Wilcoxon Rank Sum Test, **** adjusted P < 0.0001; ns=no significance. **(b)** Representative images of immunostaining against FABP4 and 5 showed that FABP4 and 5 were upregulated in the lung ECs of *Egln1^Tie2Cre^* mice compared to WT mice. Lung tissues were co-stained with anti-FABP4 or anti-FABP5 and anti-CD31 (marker for ECs). Nuclei were counterstained with DAPI. FABP4 and 5 expression was upregulated in pulmonary vascular ECs in CKO mice. V, vessel. More than 3 lungs from each group of mice were checked. (**c**) Representative Western Blotting demonstrating upregulation of FABP4 and 5 protein expression in the lung lysate isolated from CKO lungs compared with WT mice. (**d**) Representative Western Blotting showing an increase of FABP4 and 5 protein expression in the lung lysate isolated from Sugen5416/hypoxia (SuHx) -exposed lungs or monocrotaline (MCT) -exposed lungs compared with basal rats. (**e, f**) QRT-PCR analysis and Western Blotting demonstrating an upregulation of FABP4 and 5 in the isolated PAECs from IPAH patients compared with healthy donor lungs. Each sample represents PAECs from individual patients or healthy controls (n=3). t test (**c, d, e**). *P<0.05; **P<0.01; Scale bar, 50 μm.

### Plasma FABP4/5 levels were upregulated in PAH patients

We previously employed plasma proteome analysis in patients with PAH using a high-throughput multiplex proximity extension assay (PEA) using Canada cohorts^32^. We found that plasma FABP4 and FABP5 protein levels were upregulated in patients with PAH (**Fig 2a and 2b**). FABP4 protein was positively correlated with FABP5 protein level in PAH samples (**Fig 2c**). Comparison of FABP4 and FABP5 levels with hemodynamics and biochemical parameters in the human subjects showed that levels of FABP5 were positively correlated with mean pulmonary arterial pressure (mPAP), pulmonary vascular resistance (PVR) and NT-proBNP, the latter of which is a well-established and powerful marker of heart failure risk. The levels of FABP4 were also positively correlated with PVR, and NT-proBNP, but not mPAP (**Fig 2d-2f**). The levels of FABP4 and FABP5 were negatively correlated with the cardiac index, stroke volume, and estimated glomerular filtration rate (eGFR) (**Fig 2g-2i**). These data suggest that plasma FABP4 and FABP5 were elevated in the PAH patients.

**Figure 2.**
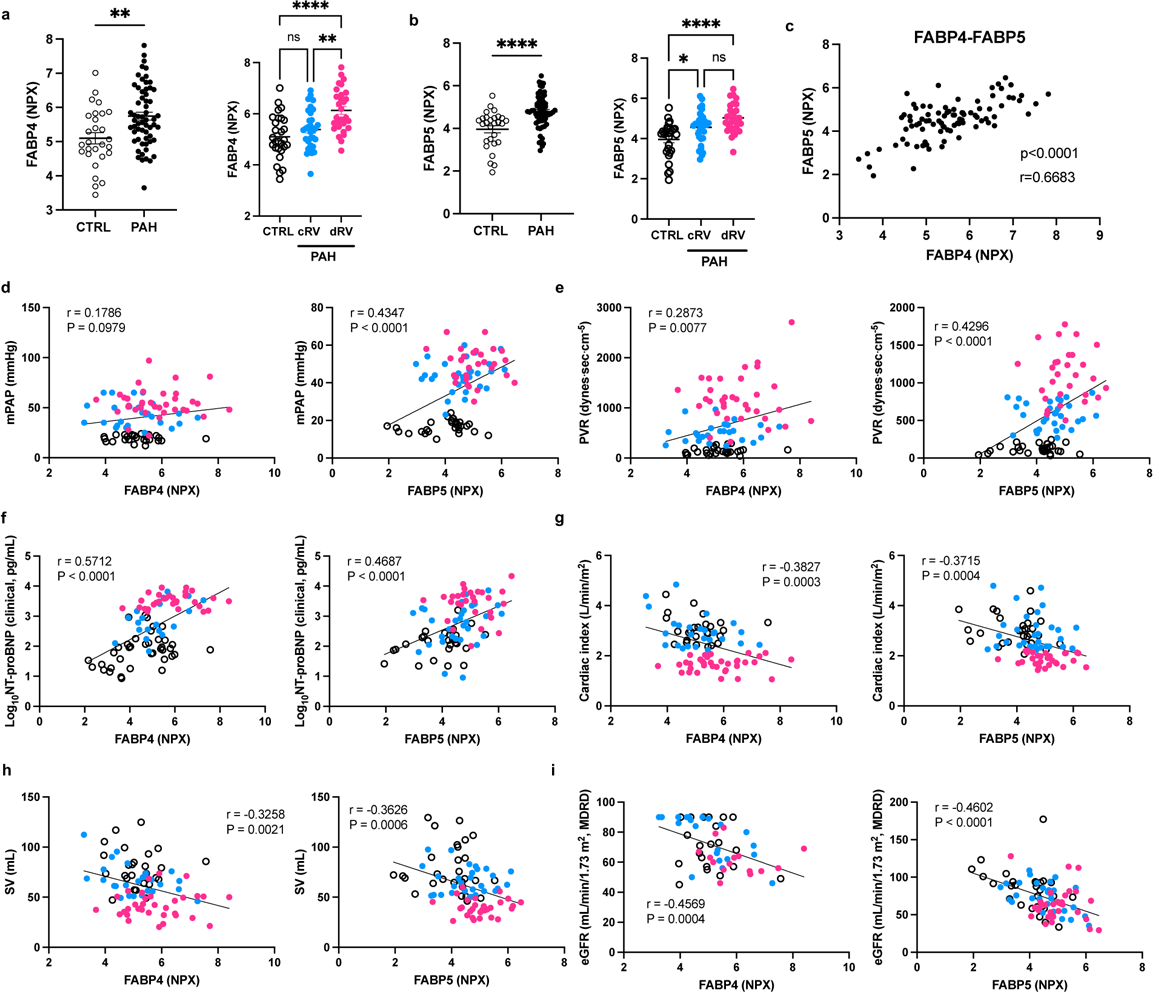
Upregulation of the plasma FABP4 and 5 in patients with PAH. (**a, b**) Plasma protein NPX levels of the FABP4 (**a**) and 5 (**b)** in donors (CTRL) and patients with PAH from Canada cohorts. Scatter dot plots show individual values (n). According to contemporary Cardiac Index (CI) measured by right heart catheterization at the time the blood sample was drawn. Patients with PAH were categorized as compensated RV (cRV, CI> 2.2 L/min/m^2^) and decompensated RV (dRV, CI ≤2.2 L/min/m^2^). CTRL N=28, cRV N= 31, dRV N=29. (**c**) Plasma protein NPX level of FABP4 were correlated with plasma protein NPX level of FABP5 in the human subjects including both donors and PAH patients. (**d-i**) Pearson’s correlation coefficient of the levels of FABP4 and 5 and hemodynamics/biomedical parameters of PAH patients with associated P value is shown in each graph. mPAP, mean pulmonary arterial pressure; PVR, pulmonary vascular resistance; SV, strove volume; eGFR, estimating glomerular filtration rate. t test (left panels of **a, b**). ANOVA followed by Turkey post hoc analysis was used for statistical analysis (right panels of **a, b**). Pearson’s correlation test (**c-i**). **P<0.01; ****P<0.0001. ns=not significant.

### *Fabp4/5* deletion protected *Egln1* deficiency-induced PH in mice

To gain insight into the role of FABP4/5 in the development of PAH, we obtained a mouse model with *Fabp4* and *Fabp5* double knockout in the whole-body (*Fabp4/5^-/-^*) from Gökhan S. Hotamisligil, Harvard School of Public Health^33^. We first exposed *Fabp4/5^-/-^* mice to chronic hypoxia for 3 weeks. We found that *Fabp4/5^-/-^* mice were protected from hypoxia-induced PH development (**Extended Data Fig 1**). Next, we generated *Egln1^Tie2Cre^*/*Fabp4/5^-/-^* (TKO) mice by breeding *Fabp4/5^-/-^*mice with *Egln1^Tie2Cre^* mice (**Fig 3a**). Western blotting confirmed that FABP4/5 proteins were undetectable in lungs of TKO mice but increased in CKO mice compared to WT controls (**Fig 3b**). As CKO mice develop spontaneously progressive PH^27^, we then evaluated the PH phenotype in TKO mice. Age and gender-matched TKO mice were compared with control *Egln1^f/f^*/*Fabp4-5^-/-^*(DKO) mice at the ages of between 2∼3.5 months. CKO mice showed increases in RVSP and right heart vs left heart and septum ratio (indicative of RV hypertrophy), whereas TKO mice exhibited a reduction of RVSP and RV hypertrophy compared to CKO mice (**Fig 3c and 3d**). These data indicated that FABP4/5 contribute to the pathogenesis of severe PH in *Egln1*-deficient mice.

**Figure 3.**
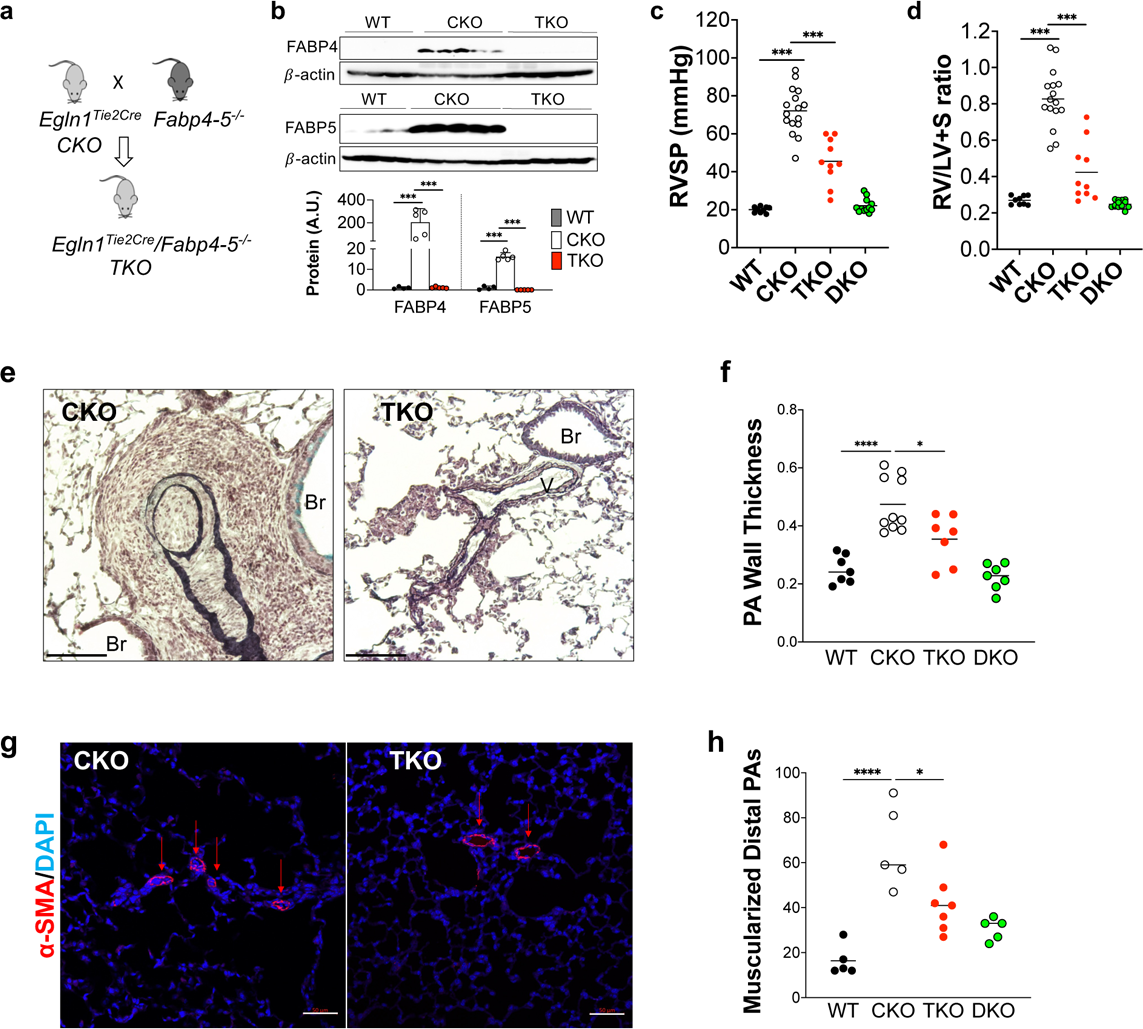
FABP4 and 5 deletions protected from *Egln1*-deficiency-induced PH in mice. **(a)** A diagram showing the strategy for generation of FABP4 and 5 double knockouts in *Egln1^Tie2Cre^* mice (TKO). **(b)** Representative Western blotting demonstrating diminished FABP4 and 5 protein expression in lung lysate isolated from TKO lungs compared with CKO lungs. (**c**) Dramatic reduction in RVSP was seen in TKO mice compared with CKO mice. Bars represent the mean. (**d**) Marked inhibition of RV hypertrophy in TKO mice compared with CKO mice. (**e, f**) Representative images from Russel-Movat pentachrome staining and quantification of wall thickness of PA exhibited reduced thickness in the intima, media, and adventitia layers of tissue of TKO mice compared with CKO mice. Occlusive lesions were diminished in TKO mice compared with CKO mice. Br, bronchus; V, vessel. (**g**) Exemplary images showed a reduction in muscularization of distal pulmonary vessels in TKO mice. Lung sections were subjected to immunostaining using anti-smooth muscle α-actin (SMA). (**h**) Muscularization quantification was performed by tallying SMA-positive vessels across 40 fields (at ×20 magnification) within every lung section. The results were presented as the mean value along with the standard deviation (mean ± SD) based on a sample size of 5 to 10 mice per group. ANOVA followed by Turkey post hoc analysis was used for statistical analysis (**b, c, d, f** and **h**). *P<0.05; ***P<0.001, and ****P<0.0001.

To further determine whether FABP4/5 regulate pulmonary vascular remodeling in CKO mice, we performed Russel-Movat pentachrome staining and immunostaining of lung tissues for α-smooth muscle actin (SMA)^10,11,22^. Examination of lung pathology showed that in comparison to CKO mice, TKO mice exhibited a marked reduction of pulmonary wall thickness and distal pulmonary arterial muscularization as assessed by α-SMA immunostaining (**Fig 3e-3h**). Taken together, genetic deletion of *Fabp4/5* attenuated pulmonary vascular remodeling and PH in *Egln1*-deficient mice.

### Genetic deletion of *Fabp4/5* improved right heart function in PH mice

Patients with severe PAH die of right heart hypertrophy and failure. We found that plasma level of FABP4 was increased in decompensated RV failure, whereas FABP5 level was elevated in both compensated and decompensated RV failure in human PAH patients (**Fig 2a** and **2b**). Therefore, we tested if genetic *Fabp4/5* deletion will prevent right heart failure in CKO mice. Echocardiography measurements showed that RV wall thickness and RV chamber size were decreased in TKO mice compared to CKO mice (**Fig 4a**). RV contractility, assessed by RV fraction area change, was also improved in TKO mice compared to CKO mice (**Fig 4b**), suggesting that RV function was better in CKO mice after deletion of *Fabp4/5*. In addition, pulmonary arterial function, determined by PA AT/ET ratio, was improved after *Fabp4/5* deletion (**Fig 4c**). We did not observe any significant alterations in heart rate, cardiac output, left ventricular fractional shortening among these mice (**Fig 4d-4f**). These data demonstrated that FABP4/5 contributed to right heart dysfunction in PH induced by *Egln1* deficiency.

**Figure 4.**
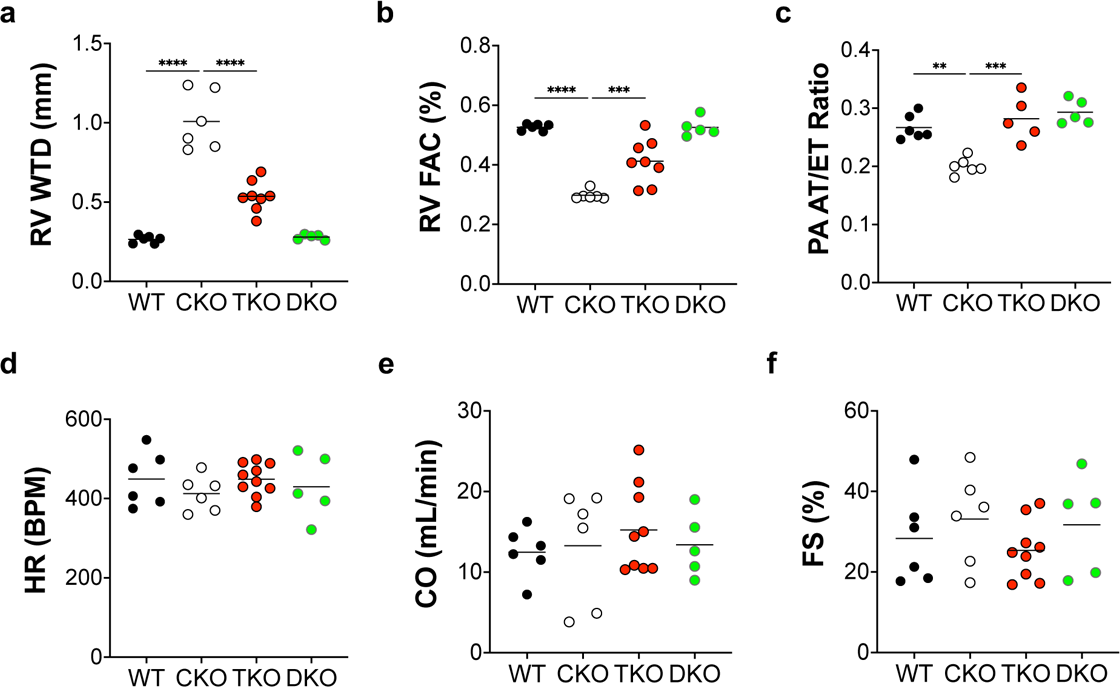
Genetical deletion of Fabp4-5 improved right heart function in PH mice. **(a)** Representative echocardiography images revealed a reduction of RV wall thickness during diastole (RVWTD) in TKO mice compared with CKO mice. (**b**) Improved RV fraction area change (RVFAC), indicating enhancing RV contractility, in TKO mice compared with CKO mice. (**c**) An increased ratio of pulmonary artery acceleration time to ejection time (PA AT/ET) in TKO mice compared with CKO mice. (**d-f**) There is no significant change of heart rate, cardiac output and left ventricular fractional shorting. ANOVA followed by Turkey post hoc analysis was used for statistical analysis (**a, b, c**). Significance levels were denoted as **P<0.01, ***P<0.001, and ****P<0.0001.

### *Fabp4/5* deletion normalized the altered expression of PH-causing and glycolytic genes

To delineate the molecular mechanisms mediated by FABP4/5 in PH development, we performed whole-transcriptome RNA sequencing analysis. Principal component analysis and the representative heatmap demonstrated that *Fabp4/5* deletion altered the overall gene expression signature in CKO lungs toward gene signature in WT lungs (**Fig 5a and 5b**). We found that 2,727 genes (adjusted p < 0.05) were significantly altered in CKO lungs compared with WT lungs (**Fig 5c** and **Extended Data Fig 2a**). Among these genes, there were 1,455 upregulated and 1,272 downregulated genes in CKO lungs. 63% of the dysregulated genes were normalized when *Fabp4/5* were deleted (**Fig 5C and Extended Data Fig 2b and 2c**). Next, we performed the enriched Kyoto Encyclopedia of Genes and Genomes (KEGG) pathways and Gene Set Enrichment Analysis (GESA) analysis based on the list of genes upregulated in CKO lungs and downregulated in TKO lungs. The KEGG analysis revealed that Epithelial mesenchymal transition, Coagulation, Hypoxia, Inflammatory response, Glycolysis, and Complement pathways were positively regulated by FABP4/5 (**Fig 5d**). All these pathways have been shown to mediate PH development. The GESA analysis showed that the enrichment of Hypoxia, and glycolysis pathways were positively regulated by FABP4/5 (**Fig 5e**). We also observed that expression levels of many genes related to PH development (*Bmpr2, Cav1, Cxcl12, Pdgfb*) and glycolysis (*Eno1, Gapdh, Ldha*) were dysregulated in CKO lungs but normalized in TKO lungs. (**Fig 5f**). qRT-PCR and western blotting were performed to confirm that mRNAs and proteins of PH and glycolysis-associated genes were altered in CKO lungs but normalized in TKO lungs (**Fig 5g and 5h**). Taken together, these data demonstrate that genetic deletion of *Fabp4/5* altered PH-associated gene expression in *Egln1*-deficient mice.

**Figure 5.**
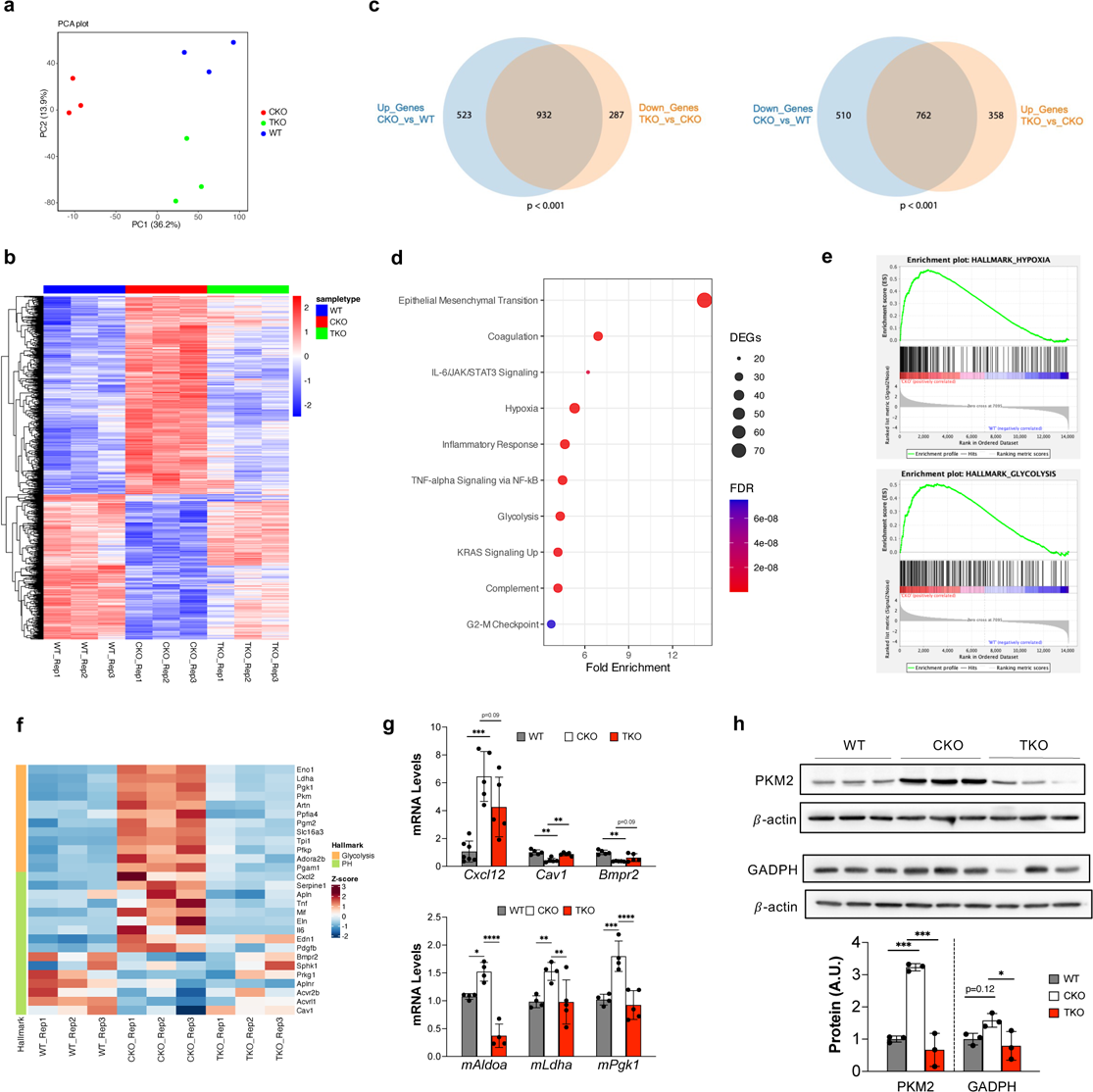
FABP4-5 regulated genes related to PH and glycolysis. **(a)** Principal component analysis showed that FABP4-5 deletion normalized the altered gene signature by *Egln1* deficiency in mice. (**b**) A representative heatmap showed that Fabp4-5 deletion altered the overall gene signature of CKO lungs toward to WT lungs. (**c**) Venn diagrams indicated FABP4-5 controlled the expression of the dysregulated genes in CKO mice. (**d**) The Kyoto Encyclopedia of Genes and Genomes (KEGG) pathway analysis of the genes upregulated in CKO lungs and normalized by FABP4-5 deletion. (**e**) Gene Set Enrichment Analysis (GESA) analysis showed the enrichment of Hypoxia and Glycolysis pathways based on the gene sets regulated by FABP4/5. (**f**) A panel of representative genes related to PH pathogenesis and glycolysis were normalized by FABP4-5 deletion in CKO mice. (**g**) qPCR analysis confirmed the PH causing genes and glycolysis related genes were altered in CKO lungs and normalized in the TKO mice by RNA-seq data. (**h**) Western blotting showing FABP4-5 deletion reduced the protein level of glycolytic genes in CKO mice. ANOVA followed by Turkey post hoc analysis was used for statistical analysis (**g** and **h**). *P<0.05; **P<0.01; ***P<0.001, and ****P<0.0001.

### scRNA Transcriptomic analysis of the ECs regulated by FABP4/5 in PH

To gain an understanding of cellular and molecular mechanisms of FABP4-5 upregulation as a contributing factor to pathogenesis of severe PH, we performed whole lung scRNA-seq analysis of WT, CKO, TKO at the age of between 2 to 3.5 months. Our data showed that ECs were one of the major cell types with significant changes in CKO mice compared to WT mice^29^ (**Fig 6a** and **6b**, and **Extended Data Fig 3a-3c**). To determine the molecular mechanisms regulated by FABP4/5 in pulmonary ECs, we performed a focused transcriptomic analysis of ECs from the scRNA-seq data. We found that 58.6% of the upregulated genes and 67.5% of the downregulated genes in the CKO mice were normalized by TKO ECs. **(Fig 6c**). Similar to the bulk RNA-seq analysis, pathway enrichment analysis on the ECs transcriptomes showed that Hypoxia, Epithelial mesenchymal transition, Glycolysis pathways were enriched in the upregulated DEGs in the CKO ECs and normalized in TKO ECs (**Fig 6d**). Expression of many ECs genes related to glycolysis (including *Eno1*, *Pgk1*, *Tpi1*, *Pkm*, *Ldha*, *Pfkl*) were upregulated in pulmonary ECs of CKO mice, but the expression of these genes was normalized in TKO mice (**Fig 6e**). scRNA-seq data showed that the alteration of glycolysis score was increased in ECs but also detectable in other pulmonary cell types such as macrophages, pericytes and T cells (**Fig 6f and Extended Data Fig 4**).

**Figure 6.**
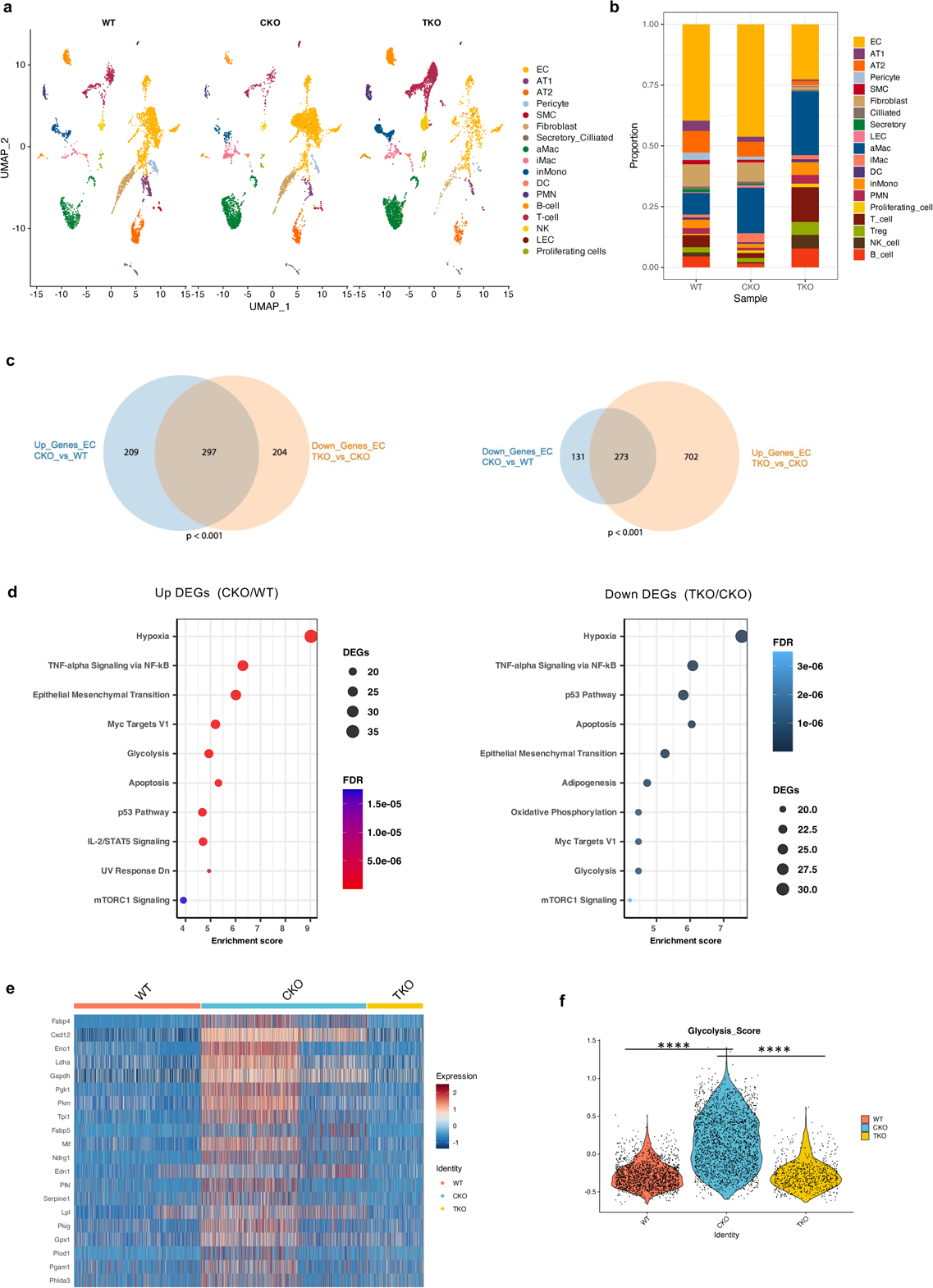
Single-cell transcriptomics analysis on FABP4-5 regulated PH development. **(a, b)** ScRNA-seq analysis demonstrated that FABP4-5 deletion partially normalized the alteration of EC subpopulations in CKO mice. **(c)** Venn diagrams indicated FABP4-5 controlled the expression of the dysregulated genes in ECs. (**d**) The Kyoto Encyclopedia of Genes and Genomes (KEGG) pathway analysis of the genes upregulated in CKO ECs and normalized by FABP4-5 deletion in ECs. (**e**). A heatmap analysis based on the scRNA-seq analysis on ECs population showed that the glycolytic genes were upregulated in CKO lungs and normalized in TKO ECs. (**f**) The glycolytic score calculation showed that glycolysis is controlled by FABP4-5 in ECs.

### Single-cell and spatial transcriptomics analysis identified distal arterialization was regulated by FABP4/5

Recent studies employing scRNA-seq analysis demonstrated that lung ECs contained arterial (expressing markers *Cxcl12, Sparcl1, Edn1, Hey1*)^34^, venous (*Prss23*, *Vwf*)^34^, general capillary ECs (*Gpihbp1*, *Plvap1*), and aerocytes (*Car4, Ednrb*)^34,35^. Our recent studies demonstrated that general capillary EC derived arterial ECs were accumulated in the distal lung of PH mice and human IPAH patients^24^. Using similar markers to define EC subpopulations (**Extended Data Fig 5**), our scRNA-seq data showed remarkable decreases in both general capillary EC and aerocyte proportion, but an increase in arterial EC proportion in CKO mice compared to WT mice, and the proportion of arterial ECs and capillary ECs were normalized in TKO mice compared to CKO mice (**Fig 7a-7c**). We also performed Visium spatial transcriptomic analysis using Visium platform and observed the reduction in arterial ECs and increase in capillary ECs in the distal lung in the TKO mice compared to CKO mice (**Fig 7d and Extended Data Fig 6**). Both gCap marker Ghibp1 and aCap marker Car4 were partially restored in the TKO lungs compared to CKO lungs (**Fig 7e**). We further profiling the arterial markers and confirm that arterial EC markers including *Sox17, Tm4sf1, Cxcl12, Sparc* were reduced in the ECs of TKO lungs compared to CKO lungs by single-cell transcriptomics analysis (**Fig 7f**). Similar data was also observed in the whole lung of the bulk RNAseq analysis (**Fig 7g**). The classical arterial marker SOX17 were further validated by western blotting on the lung lysates and immunostaining analysis at the distal microvascular bed. Taken together, our data suggest that endothelial distal arterializaiton and glycolytic programming induced by FABP4/5 contributes to pathogenesis of severe PH in *Egln1*-deficient mice.

**Figure 7.**
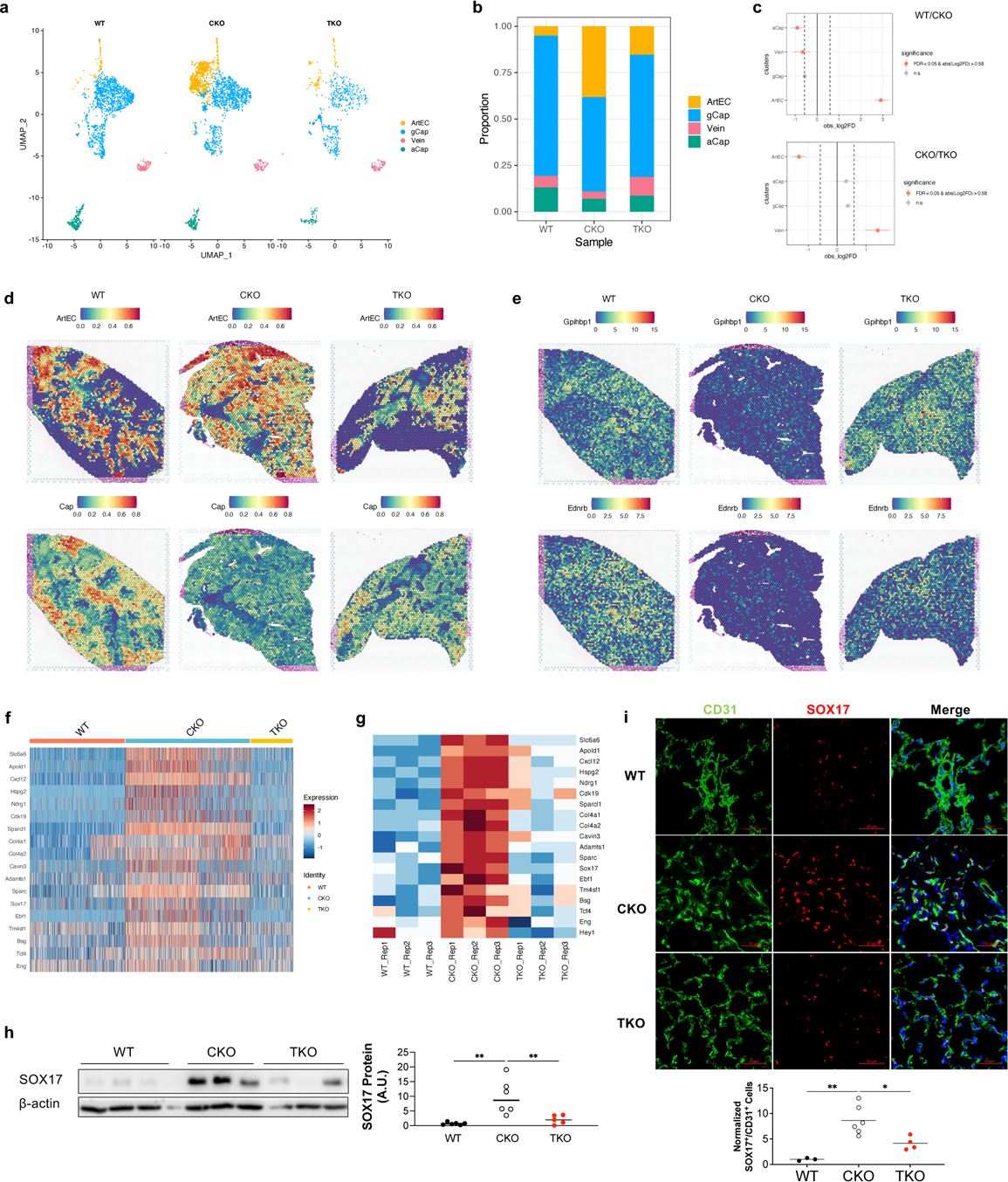
Single-cell and spatial transcriptomics analysis identified induction of distal arterialization by FABP4-5 in PH. **(a, b)** A representative UMAP and a cellular proportion plot showing the EC subpopulation change between groups. Upregulation of arterial EC proportion in CKO lungs was inhibited in TKO lungs. (**c**) Statistical analysis showed that an increase in arterial ECs and a reduction of aCap in CKO lungs were rescued in TKO lungs. (**d**) Integration of scRNAseq and Visium spatial data revealed increased arterial ECs and decreased gCap ECs in the distal capillary bed of CKO mice, which was normalized in the TKO mice. The same WT and CKO lungs were used in our previous publication^24^. The visualization shows the predicted spatial distribution of arterial ECs and capillary ECs within the lungs. (e) A spatial Plot showed that both gCap marker Gpihbpi and aCap marker Car4 were reduced in the CKO lungs and restored in the TKO lungs. (**f**) A heatmap showing a panel of representative genes related to arterial EC markers were increased in the CKO ECs and normalized in TKO ECs based on the scRNAseq data. (**g**) A heatmap based on the Bulk RNAseq data showing the rescue of arterial gene programming in TKO lungs compared to CKO lungs. (h) Western blotting demonstrated the classical arterial marker SOX17 was increased in the CKO lungs and inhibited in the TKO lungs. (i) Immunostaining against SOX17 demonstrated the increase in distal arterial ECs in CKO lungs, which was rescued in the TKO lungs. ANOVA followed by Turkey post hoc analysis was used for statistical analysis (**h** and **i**). *P<0.05; **P<0.01.

### FABP4/5 induced ECs dysfunction in PH

To understand the direct impact of FABP4/5 on PAEC glycolysis *in vitro*, we overexpressed FABP4/5 in hPAECs using lentivirus-mediated gene overexpression (**Fig 8a**) and performed glycolytic function analysis on Seahorse XF equipment. FABP4/5 overexpression stimulated glycolysis and increased cell proliferation in hPAECs (**Fig 8b and 8c**). Glycolysis has been demonstrated as an important mechanism in the development of PAH^20–23^. Pulmonary vascular ECs sustain the proliferative and anti-apoptotic phenotypes that are depended on glycolysis. To determine whether upregulation of glycolysis by FABP4/5 overexpression contributes to EC proliferation, we assessed EC proliferation (BrdU incorporation assay) in the presence of the glycolysis inhibitor, 2-DG (5 μM). Our data showed that 2-DG treatment significantly inhibited FABP4/5-induced PAEC proliferation (**Fig. 8d**). Consistent with the glycolytic status, TKO mice exhibited a reduction in EC proliferation in the lung tissue compared to CKO mice (**Fig. 8e**).

**Figure 8.**
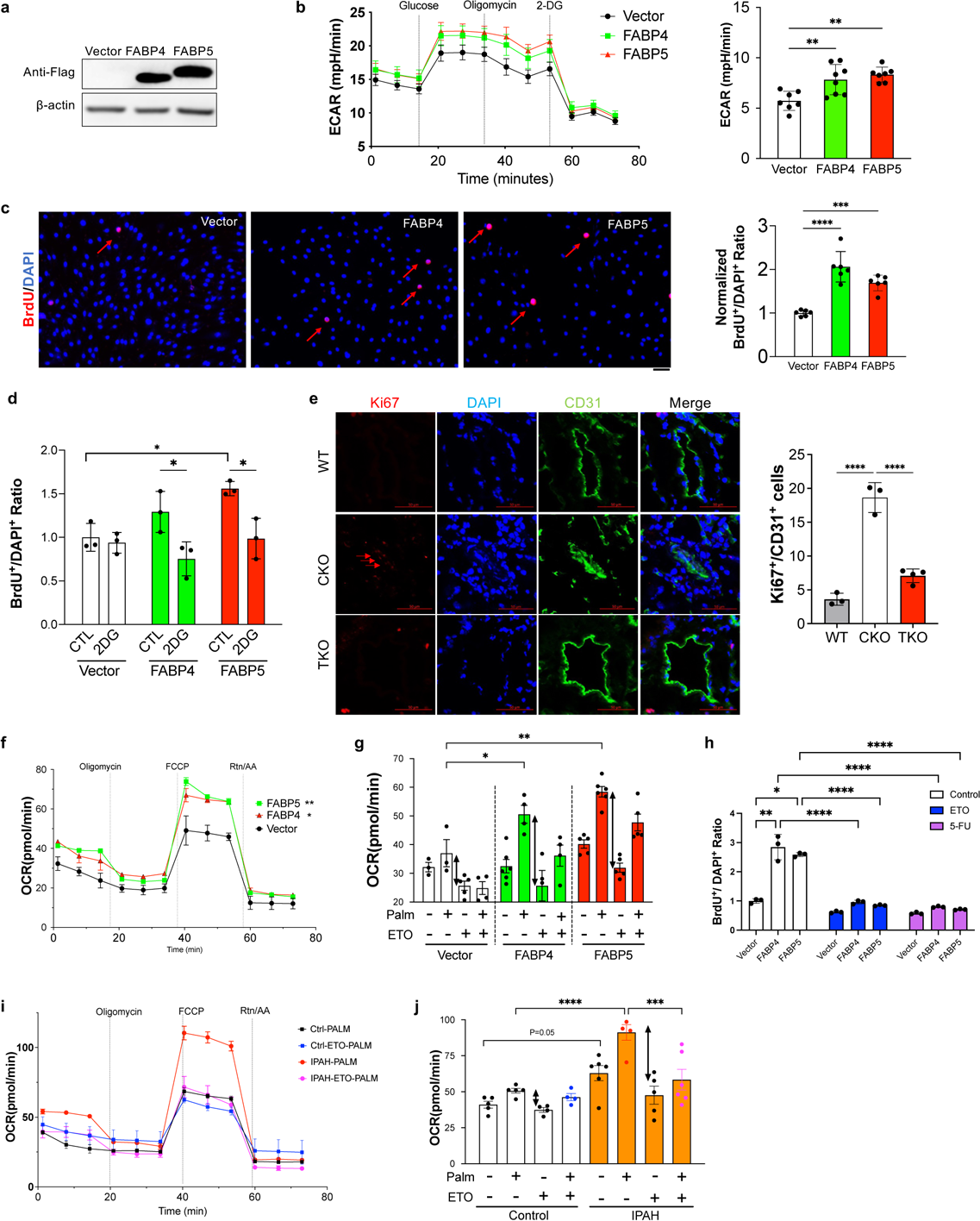
Overexpression of FABP4-5 induced ECs dysfunction. **(A)** Overexpression of FABP4 and 5 fused with Flag tag in hPAECs using lentiviruses mediated gene overexpression. (**B**, **C**) FABP4-5 overexpression promoted ECs glycolysis and proliferation assessed by BrdU incorporation assay. (**D**) Glycolytic inhibitor 2-DG treatment inhibited FABP4-5 induced PAEC proliferation. (**E**) FABP4-5 deletion in vivo reduced lung PAECs proliferation in TKO mice compared with CKO mice. (**F** and **G**) Overexpression of FABP4-5 promoted fatty acid oxidation (FAO) in hPAECs in the presence of palmitic acid. The double arrows label the OCR generation contributed by FAO. (**H**) Both CPT1α inhibitor Etomoxir (ETO, 20 µM) and DNA synthesis inhibitor (5-FU, 20 µM) blocked FABP4-5 induced ECs proliferation. (**I** and **J**) A representative seahorse data showing upregulation of FAO in PAECs isolated from IPAH patients compared to control donors. Six lines of control or IPAH patients PAECs were tested. (**C, D, H**) For BrdU assay, each dot represents a biological replicate. The experiments were performed at least three times. Student t test (**D**). ANOVA followed by Turkey post hoc analysis was used for statistical analysis (**B, C, E, F, G, H, J**). Significance levels were denoted as *P<0.05, **P<0.01, ***P<0.001, and ****P<0.0001.

The classical function of FABP4/5 is related to fatty acid binding for intracellular delivery into intracellular compartments, including mitochondria for fatty acid oxidation (FAO)^36^. Endothelial loss of CPT1α, a rate-limited enzyme for FAO, has been shown to cause impaired vascular sprouting via blocking *de novo* nucleotide synthesis required for DNA replication^37^. Our data showed that overexpression of FABP4 or FABP5 in hPAECs increased mitochondrial respiration in the presence of palmitate (the salts and esters of palmitic acid), which was blocked by CPT1α inhibitor Etomoxir (ETO)^38^, indicating that FABP4/5 promoted FAO in hPAECs (**Fig 8f and 8g**). To further determine whether FAO and FAO-derived dNTPs are required for FABP4/5 induced ECs proliferation, we treated FABP4/5 overexpressing hPAECs with ETO and 5-FU (DNA synthesis inhibitor). Our data showed that both inhibitors reduced FABP4/5-induced ECs proliferation (**Fig 8h**). Furthermore, we compared the FAO capacity between control and IPAH PAECs using Seahorse FAO assay. Our data showed that 5 of 6 lines of PAECs from IPAH exhibited upregulation of FAO compared to control PAECs (**Fig 8i and 8j**), consistent with previous publication that upregulation of FAO in lungs of PAH patients^39^.

Recent studies showed that the neointima in PH is derived from PASMCs using a house dust mite mouse model^40^, and our previous studies demonstrated that PAH ECs promoted PASMC proliferation via paracrine effect^28^. Thus, we co-cultured FABP4/5-overexpressing hPAECs with human PASMCs and found that FABP4/5-overexpressing ECs promoted PASMCs proliferation evident by increased BrdU incorporation measuring DNA replication **(Supplemental Fig 7**).

### HIF-2α mediates glycolysis induced by FABP4/5 in pulmonary ECs

Multiple transcriptional factors such as FOXO1, FOXM1, c-MYC, KLF2, HIF-1/2α, YAP1, and Notch are involved in activation of metabolism in pulmonary ECs^41,42^. To define the molecular mechanisms responsible for FABP4/5-mediated activation of EC glycolysis, we performed the iRegulon analysis (a method to detect transcriptional factors, targets and motifs from a set of genes)^43^ based on the overlapping genes upregulated (>2 fold) in CKO mice and normalized in TKO mice. We found that both HIF-2α and HIF-1α were enriched based on published literature and iRegulon analysis (**Fig 9a).** Western Blotting analysis confirmed that HIF-2α protein expression was upregulated in CKO lungs and normalized in TKO lungs (**Fig 9b**). HIF-1α was not considered for further investigations because our previous study showed that HIF-1α deletion in CKO mice did not protect from severe PH^27^; and therefore, it is unlikely that HIF-1α is involved in FABP4/5-mediated PH pathogenesis. To reduce the risk of excluding potential factors in addition to HIF-2α, we also validated p53 and c-Myc expression. Our data showed that neither p53 nor c-Myc was upregulated in CKO lungs and normalized in TKO lungs (**Fig 9b**). Based on RNA-seq data from WT, CKO mice, and *Egln1^Tie2Cre^/Hif2a^Tie2Cre^* (EH2) double knockout mice^27^, glycolysis-related genes (including *Eno1*, *Ldha*, *Pkm2*) were highly induced in the CKO lungs and normalized in *EH2* lungs (**Fig. 9c**), suggesting that HIF-2α is a key factor which regulates glycolysis downstream of FABP4/5 in CKO mice. To determine whether HIF-2α is involved in FABP4/5-induced cell proliferation, we knocked down HIF-2α in FABP4-5-overexpressing PAECs using HIF-2α specific siRNA. HIF-2α inhibition reduced FABP4/5-induced EC proliferation (**Fig 9d**).

**Figure 9.**
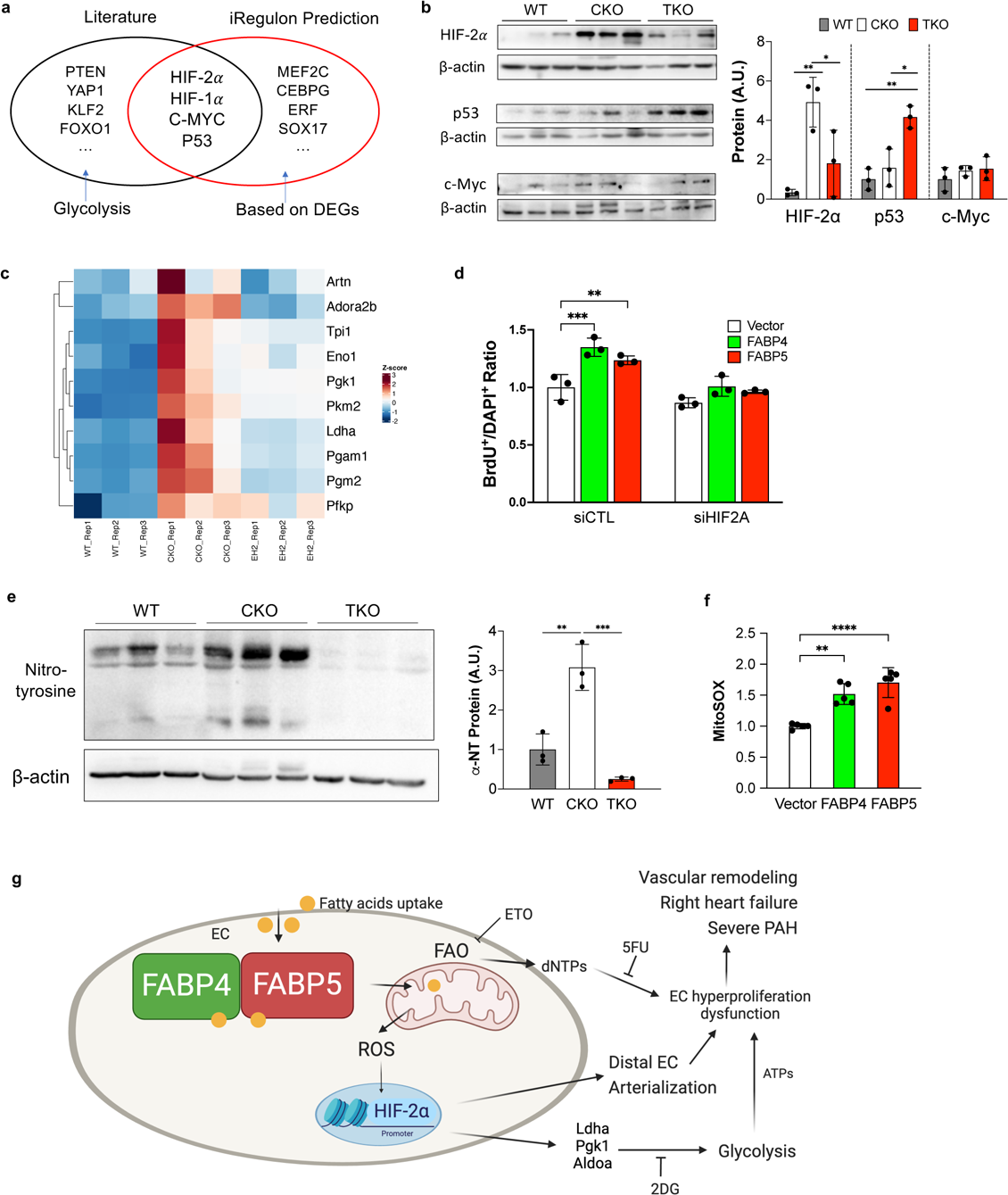
HIF-2α mediated glycolysis induced by FABP4-5 in pulmonary ECs. **(A)** A diagram showing the predicted transcription factors based on the DEGs and literature. (**B**) Western blotting demonstrated HIF-2α but not p53 or C-Myc was upregulated in CKO lungs and normalized in TKO lungs. (**C**) A representative heatmap showed that glycolytic genes were depended on HIF-2α using WT, CKO mice, and *Egln1^Tie2Cre^/Hif2a^Tie2Cre^*(EH2) mice. (**D**) HIF-2α knockdown inhibited FABP4-5 induced endothelial proliferation. (**E**) Nitrative stress assessed by protein nitrotyrosine modification was reduced in TKO lungs compared to CKO lungs. (**F**) Overexpression of FABP4 and 5 promoted mitochondrial ROS levels in HPAECs. (**G**) A diagram showing our proposed model. Our study addresses a novel role of lung endothelial FABP4-5 controlling PAECs accumulation through increased glycolysis in the pathogenesis of PAH. By facilitating fatty acid uptake and translocation into mitochondria, FABP4-5 promote FAO and ROS generation, which activates HIF-2α signaling to promote endothelial glycolysis. For BrdU assay in (**D**) and mitochondrial ROS assay in (**F**), each dot represents a biological replicate. The experiments were performed at least three times. ANOVA followed by Turkey post hoc analysis was used for statistical analysis (**B, D, E, F**). Significance levels were denoted as *P<0.05, **P<0.01, ***P<0.001, and ****P<0.0001.

Since FAO increased mitochondrial ROS levels which promotes stabilization of HIF-α through prolyl hydroxylases inhibition^44,45^, we hypothesize that FABP4/5 upregulate HIF-2α via FAO/mitochondrial ROS generation in PAECs. To test this hypothesis, we first performed staining against oxidative/nitrative stress markers^46^ such as nitrotyrosine^47^ in the lungs from WT, CKO, and TKO mice and found that *Fabp4/5* deletion reduced nitrotyrosine expression (**Fig 9e**). Next, we measured mitochondrial ROS levels using MitoSOX Red Mitochondrial Superoxide Indicator^48^ in FABP4/5-overexpressing hPAECs and found that FABP4/5 overexpression upregulated the mitochondrial ROS levels (**Fig 9f**). Taken together, these studies demonstrate that ROS/HIF-2α signaling is required for FABP4/5 to induce ECs dysfunction in PH.

## Discussion

In this study, we demonstrated that FABP4 and FABP5 were markedly upregulated in pulmonary ECs of mice with severe PH, lung tissue of IPAH patients, human PAECs, and in lung tissues of rodent PH models, including *Egln1^Tie2Cre^* mice, MCT-treated rats, and SuHx-treated rats. Plasma levels of FABP4 and 5 were also elevated in patients with PAH and correlated with hemodynamics and biochemical parameters in PAH patients. Genetic deletion of *Fabp4* and *Fabp5* protected from severe PH and right heart failure in *Egln1^Tie2Cre^* mice, supporting the critical role of FABP4/5 in pathogenesis of PH. We also observed that FABP4/5 promoted EC glycolysis and distal EC arterialization, and regulated ECs proliferation in HIF-2α-dependent manner. Deletion of *Hif2a* in ECs blocked glycolysis and ECs proliferation. Furthermore, we demonstrated FABP4/5 induced FAO and ROS generation in pulmonary ECs, which in turn contributes to HIF-2α activation (**Fig 9g**).

We provide evidence that FABP4/5 were upregulated in the lung ECs and tissue in PH rodents and IPAH patients. Overexpression of FABP4/5 was also identified in the systemic circulation of PAH patients. Although an increase in FABP4 in circulation in PAH patients was previously identified^18^, and serum FABP4 concentrations were significantly higher in patients with HF than those in non-HF subjects^49^, our study is the first report to link circulating levels of FABP4 and 5 to hemodynamics and RV failure in patients with PAH. We also determined the causative role of FABP4/5 in PH using genetic mouse models and identified the underlying molecular mechanisms whereby FABP4/5 regulate lung remodeling in PH. Circulating FABPs are seen as markers of cell injury or death. During PAH development, endothelial injury is accompanied by tissue hypoxia and inflammation. Both HIF-1α and HIF-2α expression is upregulated in PAH ECs.^26,50,51^ Previous studies also showed that FABP4 was induced during hypoxia treatment in hepatocytes in HIF-1α-dependent manner during liver ischemia/reperfusion^52^. Our data showed marked upregulation of FABP4/5 in CKO lungs with severe PH and the normalization of PH parameters in EH2 mice *in vivo* (**Supplemental Fig 8**), suggesting that FABP4/5 are novel downstream targets of HIF-2α in PAECs. FABP4 is regulated by VEGF and contributes to EC proliferation, migration, and angiogenesis^53,54^. Hwangbo et al showed that endothelial Apelin/Apelin receptor signaling through FoxO1 regulated FABP4 but not FABP5 during the tissue fatty acid uptake in skeletal muscles^16^. Further studies are warranted to determine the upstream signaling regulating FABP4/5 in PH.

Previous studies showed that both FABP4 and FABP5 played important roles in the pathogenesis of chronic metabolic diseases, including obesity and diabetes^36^. The *Fabp4-Fabp5* double knockout mice (*Fabp4/5*^-/-^) were protected from high fat diet-induced weight gain and insulin sensitivity, as well as hepatic steatosis^55^. Accumulating evidence shows that PAH is a systemic metabolic disease^4–6^. The prevalence of obesity in PAH doubled between 2006 and 2016^56^, and the insulin resistance is likely a risk factor for PAH^6^. In the present study, we provide the first genetic evidence that *Fabp4/5* deletion attenuated PH development, including inhibition of RVSP and pulmonary vascular remodeling, as well as prevented right heart failure induced by *Egln1* deficiency. Our data are consistent with previous studies showing that FABP5/7 inhibition using SBFI-26 aggravated pulmonary arterial fibrosis and PH secondary to left heart disease induced by ligation of left main artery in mice^57^. Taken together, our data indicate that FABP4/5 contribute to the pathogenesis of PAH and right heart failure.

FABP4 and 5 are expressed in multiple PH-related cell types, including ECs and macrophages in human and mouse lungs based on LungMAP datasets (**Supplemental Fig 9**). Our data showed that *Fabp4 and 5* were mainly upregulated in ECs in CKO mice. We then demonstrated that FABP4/5 promoted endothelial dysfunction, including augmentation of arterial programming, glycolysis and proliferation, and altered paracrine effects in ECs *in vitro* and *in vivo*. In general, FABPs function as lipid chaperones to facilitate fatty acid trafficking. Increased intracellular influx of fatty acids facilitated by FABP4/5 upregulates the fatty acid oxidation in ECs, which likely provides the sources of dNTPs for DNA synthesis and ECs hyperproliferation in PH condition. This finding is consistent with previous study showing that heterozygous deletion of *Cpt1a*, the rate limited enzyme for fatty acid oxidation, protected from hypoxia-induced PH in mice^39^. Moreover, our studies identified a novel role of FABP4/5 in regulation of endothelial glycolysis through activation of ROS/HIF-2 signaling in PH. Previous studies showed that free fatty acids upregulated protein level of HIF-2α in clear cell renal cancer cells^58^. It is likely that FAO derived from FABP4/5 binding of fatty acid upregulates HIF-2α expression. Moreover, our study also found that overexpression of FABP4/5 reduced endothelial PPARγ protein levels whereas knockdown of FABP4/5 upregulated endothelial PPARγ protein but not mRNA expression (**Supplemental Fig 10**), which might further lead to EC dysfunction and PH development.

Both FABP4 and FABP5 are expressed in alveolar macrophages^59^. However, previous studies showed that FABP4 and FABP5 regulated different signaling pathways in macrophages compared to other cell types. For example, FABP4 was highly expressed in alveolar macrophages in COVID-19 patients, which could contribute to obesity-associated severity of COVID-19^60^. Another study demonstrated that loss of FABP5 promoted pro-inflammatory reprogramming of macrophages, increasing the expression of pro-inflammatory cytokines and transcription factors^61^. One might speculate that macrophage FABP4 and FABP5 might play a role in the pathogenesis of PH. To answer this question, we analyzed the macrophage populations in *Fabp4/5*-deficient mice. Our data showed that *Fabp4/5* deletion did not change the cellular distribution in macrophage subsets (**Supplemental Fig 11**), suggesting that macrophage FABP4/5 might play an opposite role in PH development. Future studies are needed to employ macrophage-specific deletion of *Fabp4/5* in mice to determine the role of macrophage FABP4 and FABP5 in PH development.

FABPs are traditionally considered cytoplasmic proteins. FABPs were identified in the circulation under obesity-associated diseases, cancer and in PAH^36^. A previous study showed that circulating FABP4 released from adipose tissue directly enhances tumor stemness and aggressiveness through activation of the IL-6/STAT3/ALDH1 pathway^62^. Exogenous FABP4 can induce human coronary artery smooth muscle cells proliferation and migration, as well as suppress eNOS expression in human umbilical vein ECs and cardiomyocyte contraction in vitro. However, the role of exogenous FABP5 is largely unknown. Future studies are needed to investigate whether blocking exogenous FABP4/5 will inhibit PH development in preclinical models.

There are several limitations in the current studies. Firstly, it remains unknown whether the upregulation of plasma FABP4 and 5 are secreted from lung ECs, other lung cell types or adipose tissue in PAH patients. Secondly, although our data point out that *Fabp4/5* are primarily upregulated in the lung ECs in PH, our study employed a global loss of *Fabp4/5* (deletion of *Fabp4/5* in all cells of the body). Therefore, the cell specific roles of FABP4 or FABP5 in PAH remain to be studied.

In summary, our studies showed that plasma levels of FABP4 and 5 were elevated in patients with PAH and positively correlated with hemodynamics, as well as inversely correlated with RV function. Our data support a pathogenic role of FABP4/5 in mediating distal EC arterialization and endothelial glycolytic programming, FAO, and pulmonary arterial remodeling in PH. Additionally, our data suggest that targeting FABP4/5 or FAO signaling may represent a novel therapeutic approach to inhibit pulmonary vascular remodeling in human PAH.

## Methods

### Human Samples

The use of archived human lung tissues and cells was granted by the UA Institutional Review Board. Human IPAH patients and healthy donors’ type 3 arterial ECs were obtained from the Pulmonary Hypertension Breakthrough Initiative. Both IPAH and healthy donors PAECs were isolated from arteries that are less than 1mm thickness according to cell isolation protocol provided by PHBI. A table summarizing clinical and demographic characteristics of IPAH patients and FD are provided in **Extended Data Table 1**. The normal human pulmonary arterial ECs (HPAECs) were purchased from Lonza (Alpharetta, GA, USA). Human plasma proteomics analysis was described previously^32^. Briefly, blood samples from 60 patients with PAH at the IUCPQ in Canada were obtained. All experimental procedures were performed with the approval of Laval University and the IUCPQ Biosafety and Ethics Committees (CER#20735). All patients provided and signed informed consent form. The characteristics of the patients were published in previous study^32^. Plasma proteomic analysis protocol was published previously^32^. Fasting blood samples were collected into EDTA tubes and analyzed using high-throughput multiplex PEA technology. Two 384-plex panels focusing on cardiometabolic, and neurology-related proteins were used. Briefly, the PEA approach uses matched pairs of antibodies conjugated to complementary DNA oligonucleotide tags to specifically bind to target proteins. Next, amplified targets were quantified by next generation sequencing, resulting in log-base-2 normalized protein expression (NPX) values.

### Mice

All the mice used in this study were C57BL/6J background. *Egln1^Tie2Cre^* (CKO) and *Egln1/Hif2a^Tie2Cre^* (EH2) mice were generated as described previously^27^. *Fabp4-5^-/-^* mice were obtained from Gökhan S. Hotamisligil, Harvard School of Public Health^33^. CKO mice were bred with *Fabp4-5^-/-^* mice to generate *Egln1^Tie2Cre^*/*Fabp4-5^-/-^* (TKO) and *Egln1^f/f^*/*Fabp4-5^-/-^* (DKO) mice. Both male and female *Egln1^f/f^* (designed as WT), CKO, TKO and DKO little mates at the age of 5 weeks to 3.5 months were used in these studies. The animal care and study protocols were reviewed and approved by the Institutional Animal Care and Use Committees of the University of Arizona.

### Rats

Male Sprague Dawley rats (Charles River Laboratories) at the age of 6 weeks (150 g to 160 g) were randomized for two groups and treated with monocrotaline (MCT) (subcutaneous, 32 mg/kg body weight) or saline and rested for 21 days. Male Sprague Dawley rats (Charles River Laboratories) at the age of 6 weeks (150 g to 160 g) were randomized for two groups and treated with Sugen5416 (subcutaneous, 20 mg/kg body weight) or vehicle and incubated in hypoxia chamber (10% O_2_) for 3 weeks, followed by incubation in room air for 3 weeks. Lung tissues were perfused with PBS and frozen.

### Hemodynamic measurement

Right ventricular systolic pressure (RVSP) was measured with a 1.4F pressure transducer catheter (Millar Instruments) and recorded with AcqKnowledge software (Biopac Systems Inc.) as described previously^27,63^. Briefly, the catheter was inserted into the right ventricle in mice under anesthesia (100 mg ketamine/5mg xylazine/kg body weight, i.p.).

### Echocardiography

Echocardiography was performed in the University of Arizona as described previously^64^. Transthoracic echocardiography was performed on a VisualSonics Vevo 3100 ultrasound machine (FujiFilm VisualSonics Inc) using an MS550D (40 MHz) transducer. The right ventricle wall thickness during diastole (RVWTD) was obtained from the parasternal short axis view at the papillary muscle level using M-mode. The RV cross-sectional area was obtained from the parasternal short axis view at the papillary muscle level using B-mode. Pulmonary artery (PA) acceleration time and PA ejection time were obtained from the parasternal short axis view at the aortic valve level using pulsed Doppler mode. The left ventricle fractional shortening (LV FS) and the cardiac output (CO) were obtained from the parasternal short axis view using M-mode.

### Immunofluorescent staining and histological staining

Mouse lung tissues were perfused with PBS, inflated with 50% OCT in PBS, and embedded in 100% OCT for cryosectioning. For immunofluorescent staining on these fresh frozen tissue, lung sections (5 μm) were fixed with 4% paraformaldehyde and blocked with 0.1% Triton X-100 and 5% normal goat serum at room temperature for 1 hour. After 3 washes with PBS, the slides were incubated with anti-FABP4 (Cell Signaling Technology, Cat#2120S, 1:100) or anti-FABP5 (Cell Signaling Technology, Cat#39926S, 1:100), anti-CD31 antibody (BD Bioscience, Cat#550274, 1:25), anti-α-SMA (Abcam, Cat#Ab5694, 1:300), anti-CD45 (BioLegend, Cat#103101, 1:100), anti-Ki67 (Abcam, Cat#ab16667, 1:25) at 4°C overnight then incubated with Alexa 594, Alexa 488 or 647-conjugated anti-rat or anti-mouse or anti-rabbit IgG (Thermo Fisher Scientific) at room temperature for 1 h. Nuclei were counterstained with DAPI mounting medium (SouthernBiotech, Birmingham, AL, USA). Quantification of α-SMA^+^ vessels, CD45^+^ cells and Ki67^+^ ECs was performed blindly.

Mouse lung tissues were perfused with PBS and fixed with 10% formalin via tracheal instillation at a constant pressure (15 cm H_2_O) and embedded in paraffin wax. Lung sections were stained with a Russel-Movat pentachrome staining kit (Cat #KTRMP, StatLab) according to the manufacturer’s protocol. For assessment of PA wall thickness, PAs from 40 images at 20X magnification were quantified blindly by Image J. Wall thickness was calculated by the distance between internal wall and external wall divided by the distance between external wall and the center of lumen. Lung sections from formalin-fixed mouse samples were dewaxed and dehydrated, followed by boiling in 10 mmol/L sodium citrate (pH 6.0) for 10 minutes for antigen retrieval. Slides were incubated with anti-α-SMA (Abcam, Cat #ab5694, 1:300) at 4°C overnight followed by Alexa 594 or 488 conjugated anti-rabbit IgG at room temperature for 1 h. Nuclei were counterstained with DAPI. Images were taken using ECHO REVOLVE microscopy (Discover Echo Inc). α-SMA^+^ vessels were quantified blindly in 40 fields per lung.

### Primary cultures of human lung vascular ECs

Primary human pulmonary arterial endothelial cells (HPAECs) were purchased from Lonza. Briefly, HPAECs were cultured in 0.2% gelatin (Sigma) precoated T75 flasks in EBM-2 medium supplemented with 10% FBS and EGM-2 MV singlequots (Lonza, #CC-3162), and incubated at 37 °C in a humidified 5% CO_2_ atmosphere. Cells between passage 4 and 7 were used for experiments. Primary pulmonary type 3 arterial ECs (PAECs) were obtained from PHBI and cultured as described above.

### siRNA-mediated gene knockdown and lentivirus mediated gene overexpression

FABP4 (Santa Cruz Biotechnology, inc. #sc-43592), FABP5 (ThermoFisher Scientific, #AM51331) or HIF-2α siRNA (Qiagen, # SI02663038) was transfected into ECs using HMVEC-L Nucleofector kit (Lonza, # VPB-1003) with Nucleofector^TM^ 2b device (Lonza, #AAB-1001) with program A-034. For lentivirus generation, human FABP4 and 5 ORFs with Flag tag in C terminal (GenScript USA, Inc.) were cloned into pLV-mCherry (Addgene, #36804) plasmids via replacing FABP4 or FABP5 with mCherry. Helper plasmids PSPAX2 vector (Addgene, #12260) and pMD2.G vector (Addgene, #12259) were co-transfected with pLV-FABP4, or 5 or pLV-mCherry expression vector using lipofectamine 3000 (ThermoFisher Scientific) into 293-T cells to generate lentivirus.

### *In vitro* cell proliferation analysis

For assessment of EC proliferation after FABP4 or 5 overexpression, HPAECs were infected with pLV-FABP4 or 5 lentivirus and seeded on coverslips to reach 40% to 60% confluency. Around 48 hrs post infection, cells were starved overnight in FBS-free medium. 2.5% FBS and 5-bromo-20-deoxyuridine (BrdU, Sigma, Cat #B5002) was added (10 µM) for another 4 hrs. Fixed cells were immunostained with anti-BrdU antibody (BD bioscience, cat#347580), followed by Alexa 594-conjugated anti-mouse IgG (Life Technology). Nuclei were counterstained with DAPI. Images were taken using REVOLVE microscopy (Discover Echo Inc.) at 4 X magnification for blind quantification.

### Glycolysis and fatty acid oxidation Seahorse assay

For glycolysis assay, HPAECs were transfected with siFABP4/5 or control siRNA or infected with lentiviruses overexpressing FABP4 or 5 or mcherry. 48 hours post-transfection, HPAECs were seeded at the density of 30,000 cells per wells in the Seahorse XF96 microplate for overnight incubation. The assay medium supplemented by 2 mM glutamine in Seahorse XF Base Medium was prewarmed in 37 °C incubator. Cell culture growth medium was replaced by prewarmed assay medium and the microplate was placed into a 37 °C non-CO_2_ incubator for 45 minutes to 1 hour prior to the assay. The assay cartridge was loaded with the compounds (final concentrations: Port A: 10 mM Glucose, Port B: 1µM oligomycin, Port C: 50 mM 2-DG) The Seahorse XF96 Analyzer was calibrated, and the assay was performed using the Seahorse XF Glycolysis Stress Test Assay protocol as suggested by the manufacturer. For the fatty acid oxidation assay, HPAECs were infected with lentivirus for overexpressing the FABP4, FABP5 and corresponding control mCherry. For the infection, the polybrene (Millipore Sigma, TR-1003-G) was applied to the medium at the final concentration of 0.8mg/ml. After 24 hours infection, replace the medium with fresh HPAECs culture medium for another 24 hours. At 48 hours post-infection, the HPAECs were seeded into the precoated Seahorse 96-well microplate at the density of 40,000 cells in each 96-well. HPAECs were treated with BSA control or Palmitate-BSA (Agilent, 102720-100) at the concentration of 10 µM. The Etomoxir (final concentration: 40 µM), Oligomycin (final concentration: 1.5µM), FCCP (final concentration: 1.6 µM) and Rotenone/Antimycin A (final concentration: 0.5/0.5 µM) were applied for this assay according to the manufacturer of the XF Long Chain Fatty Acid Oxidation Stress Test Kit (Agilent, 103672-100). The data was analyzed by the Seahorse Wave Desktop Software (2.6).

### Bulk RNA sequencing analysis and Single-cell RNA sequencing and spatial transcriptomics analysis

Total RNA was isolated from lung tissue from 2 to 3.5 months WT, CKO and TKO mice with Zymo Research Quick-RNA Miniprep Kits with DNase Ι digestion. RNA sequencing was carried out at the Novogene Corporation Inc. All samples were sequenced with an Illumina Novaseq 6000 platform with a pair end 150-bp read length. Raw sequencing data underwent quality control assessment using FastQC. Adapters and low-quality bases (Phred score < 33) were removed. Cleaned reads were aligned to mouse genome mm10 using STAR (v2.5.2). Gene expression levels were quantified from the aligned reads using htseq counts with the reference annotation from mm10.Ens.78. Identification of differentially expressed genes was performed using edgeR (v4.0.1) package in R with a significance threshold of adj.pvalue < 0.05 and logFC > 0.58. PCA plot with raw counts was generated by ggplot2 (v3.4.4) R package.

We collected whole lung tissues from mice at the age of 2 to 3 months for ScRNA-seq analysis. Equal number of single cells isolated from male and female WT (BioLegend, Cat#155831 or #155833) or CKO (BioLegend, Cat#155835 or #155837) or TKO (WT (BioLegend, Cat#155831 or #155833) mice was stained with Cell Hashing antibodies respectively, and loaded on 10X Genomics Chromium Single Cell Controller to generate barcoded single cells for construction of single cell cDNA libraries. ScRNA-seq was performed on a Hiseq 4000 with pair-end 150bp. Cell Ranger (v4.0.0) was used for demultiplexing and counting. The 10X Genomics mouse reference genome, refdata-gex-mm10, was used as reference genome. R package Seurat (v4.3.0) was used for data preprocessing and visualization. Initially, cells with less than 200 genes or complexity (Log10gene/UMI) smaller than 0.8 were removed. Genes expressed in fewer than 3 cells were discarded. Doublets were identified using a two-layer approach as described previously: First, scDblFinder (v1.16.0) was used to predict potential doublets using default settings in an automated and unbiased fashion. Doublets were additionally identified manually when expressing combinations of marker genes from different cell types. After doublet removal, single cells from WT, CKO and TKO samples were normalized with NormalizeData function and then integrated using an anchor based CCA pipeline. The first 30 principal components (PC) were used for clustering and the resolution was set at 0.5. UMAP (Uniform Manifold Approximation and Projection) was used for cluster visualization. Marker genes for each cluster were identified by FindAllMarkers function. A small cluster of cells showing no canonical marker gene expression and high mitochondrial ratio was deemed as low quality. This cluster was removed, and the rest of the cells were re-clustered, resulting in 26 unique clusters. Cell identity to each cluster were assigned based on previously published canonical marker genes with AddModuleScore function in Seurat according to the LungMAP CellCard and recent publications^35,65^. Differential gene expression analyses between WT, CKO and TKO samples for each cluster of interest were performed using the Wilcoxon Rank Sum Test algorithm with log normalized counts in the RNA assay as input. A threshold of 0.25 for log fold change, 0.05 for the adjusted P value and 0.1 for minimal fraction of cells was applied for downstream analysis. For sub-clustering of endothelial cells, clusters of interests (EC) were extracted and the first 20 of new principal components were used. The resolution was set at 0.1. The identity of EC subclusters were annotated based on previously published canonical marker genes. The Glycolysis Score was calculated using AddModuleScore function based on the average expression levels of each glycolytic genes *(“Eno1”, “Ldha”, “Pgk1”, “Pgm2”, “Ppf1a4”, “Slc16a3”, “Tpi1”, “Pgam1”, “Pkm”, “Artn”, “Adora2b”, “Pfkp”*) on single cell level, subtracted by the aggregated expression of control feature sets. All analyzed features were binned based on averaged expression, and the control features were randomly selected from each bin. GO enrichment and pathway analysis were performed using a web-based tool Enrichr (https://maayanlab.cloud/Enrichr/) and GSEA software (V4.3.2) to identify potential pathways and molecular functions affected between different genotypes and the retrieved combined score (log(p-value) * z-score) was displayed. Bubble charts and bar plots representing significantly enrichment pathways were generated by ggplot2 (v3.4.4) R package. Overlapping analysis was conducted using an online Gene List Venn Diagram tool (http://genevenn.sourceforge.net). Venn diagrams were generated by a web-based tool Venn Diagram Maker Online (https://www.meta-chart.com/venn). Heatmaps for bulk RNAseq were generated from scaled RPKM counts of indicated genes using pheatmap (v1.0.12) R package. scRNAseq heatmaps were plotted using DoHeatmap function in Seurat R package. Volcano plots of differentially expressed genes was performed using EnhancedVolcano (v1.20.0) R package. Genes with adj. p value < 0.05 and logFC > 0.6 were labelled as significantly differential expressed.

Mouse lung tissues were perfused with PBS and fixed with 10% formalin via tracheal instillation at a constant pressure (15 cm H_2_O) and embedded in paraffin wax. Lung tissues were sectioned into 5 μm sections. Tissue sections were placed within the fiducial frame or the etched frames of the Capture Area on the 10X Genomics Visium Spatial slides. Slides were then deparaffinated, decrosslinked and stained with H & E staining kit (Millipore Sigma). Images were acquired under Keyence BZ-X800E slide scanner. The mouse whole transcriptome probe panel is added to the deparaffinized, stained, and decrosslinked tissues. After hybridization, single stranded ligation products were released and then captured on the Visium slides. Probes are extended by the addition of UMI, Spatial Barcode and partial Read 1, followed by library preparation and sample indexed. The library was sequenced on a Hiseq 4000 with pair-end 150bp (Novogene). The raw sequencing data was analyzed by CellRanger 7.0 (10X Genomics) and Seurat V4. Visium data was integrated with scRNA-seq data. The cell annotation was transferred from scRNA-seq.

### QRT-PCR analysis

Total RNA was isolated from mouse and human cells with Zymo Research Quick-RNA Miniprep Kits with DNase Ι digestion. One microgram of RNA was transcribed into cDNA using the high-capacity cDNA reverse transcription kits (Applied Biosystems) according to the manufacturer’s protocol. Quantitative RT-PCR analysis was performed on an QuantStudio 3 system (Applied Biosystems) with the Power Track SYBR Green Master kit (Applied Biosystems). Target mRNA was determined using the comparative cycle threshold method of relative quantitation. Cyclophilin was used as an internal control for analysis of expression of mouse genes while 18s rRNA gene was used for human genes. The primer sequences were provided in **Extended Data Table 2**.

### Western Blot

Cells were collected in NP-40 lysis buffer supplemented with protease inhibitor cocktails (sigma). Equal amount of protein was loaded for SDS-PAGE and Western Blotting. The PVDF membranes were blotted with anti-FABP4 (Cell Signaling Technology, Cat#2120S), anti-FABP5(Cell Signaling Technology, Cat#39926S), anti-Flag (MilliporeSigma, Cat#F1804), anti-HIF-2α (Novus, Cat# NB100-122), anti-PKM2 (Cell Signaling Technology, Cat#4503), anti-GAPDH (Cell Signaling Technology, Cat#5174), anti-p53 (Proteintech, Cat# 10442-1-AP) or anti-c-Myc (Proteintech, Cat# 10828-1-AP) or anti-nitrotyrosine (MilliporeSigma, Cat#05-233) or anti-β-actin (Sigma-Aldrich, #A2228) antibodies.

## Data availability

Bulk RNAseq dataset (GSE261445) is available at GEO dataset. scRNAseq and Visium datasets after CellRanger analysis were available at Figshare (https://figshare.com/s/xxx). Scripts used for bulk RNA sequencing and single-cell RNA sequencing (scRNA-seq) analysis are available in GitHub (https://github.com/DaiZYlab/Fabp45). Other data that support the findings of this study are available from the corresponding author upon reasonable request.

## Statistical Analysis

Statistical determination was performed on Prism 9 (Graphpad Software Inc.). The statistical analysis for human plasma proteomic study was described previously^32^. Normally distributed variables were compared by Student’s t-test. Non-normally distributed variables were compared by Mann-Whitney U-test. For more than two groups, one-way ANOVA for normally distributed variables or the Kruskal-Wallis test for non-normally distributed variables was performed for comparison. The correlation analysis between circulating FABP4 and 5 and hemodynamics (mean PAP, PVR, CVI and stroke volume), NT-proBNP and eGFR were performed by Pearson’s correlation test. For other studies, two-group comparisons were compared by the unpaired 2-tailed Student t test for equal variance or the Welch t test for unequal variance. Multiple comparisons were performed by one-way ANOVA with a Tukey post hoc analysis that calculates corrected P values. P less than 0.05 indicated a statistically significant difference. All bars in dot plot figures represent mean. All bar graphs represent mean±SD.

## Disclosure

None.

## Author contributions

Z.D. conceived the experiments and interpreted the data. B.L., D.Y., S.L., Y. C., K.R., X.X., A.T., Y.C., Z.H., M.K., R.R, H.G., and Z.D. designed, performed experiments, and analyzed the data. Z.D. wrote the manuscript. K.S.K, V.P., C.C.G, V. V.K. revised the manuscript. Y.Y.Z, S.B., O.,V., and M.B.F. provided key experimental materials.

## Acknowledgements

We thank Dr. Gökhan S. Hotamisligil (Harvard School of Public Health) for providing the *Fabp4-5*^-/-^ mice. The authors thank the Pulmonary Hypertension Breakthrough Initiative for providing the Data/tissue samples. Funding for the Pulmonary Hypertension Breakthrough Initiative is provided under an NHLBI R24 grant (R24HL123767) and by the Cardiovascular Medical Research and Education Fund.

