## supplemental data for "Single-cell and Spatial Transcriptomics Identified Fatty Acid-binding Proteins Controlling Endothelial Glycolytic and Arterial Programming in Pulmonary Hypertension"

**Supplemental Figure legends**

**Extended Data Table 1. Clinical and demographic characteristics of PAH patients and failed donors**

| PAH or Failed Donor | Gender | Race | Age |
|---|---|---|---|
| IPAH1 | Female | White | 14 |
| IPAH2 | Female | White | 11 |
| IPAH3 | Female | White | 39 |
| IPAH4 | Female | White | 16 |
| IPAH5 | Male | White | 53 |
| IPAH6 | Male | White | 13 |
| IPAH7 | Male | Asian | 29 |
| IPAH8 | Male | Asian | 18 |
| IPAH9 | Female | White | 55 |
| IPAH10 | Female | White | 57 |
| Failed donor 1 | Female | White | 49 |
| Failed donor 2 | Female | White | 57 |
| Failed donor 3 | Male | White | 49 |
| Failed donor 4 | Male | White | 49 |
| Failed donor 5 | Female | White | 43 |
| Failed donor 6 | Male | White | 30 |
| Failed donor 7 | Female | Hispanic | 55 |
| Failed donor 8 | Male | Unknown | 13 |
| Failed donor 9 | Female | Asian | 34 |
| Failed donor 10 | Male | White | 21 |

**Extended Data Table 2. QRT-PCR primers**

| Gene name | Forward primer | Reverse primer |
|---|---|---|
| *mCxcl12* | CCAAGAGTACCTGGAGAAAGC | AGTTACAAAGCGCCAGAGCA |
| *mCav1* | GACCCCAAGCA TCTCAACGA | TTGGGATGCCGAAGATCGTA |
| *mBmpr2* | AAGAGCACAGAGGCCCAA TTC | CCATCTTGTGTTGACTCACCT |
| *mAldoa* | AGAAGGATGGAGCCGACTTTG | GTACAATGCCATTCTGCTGGC |
| *mLdha* | CAGGCTCCCCAGAACAAGATT | CTCATCCGCCAAGTCCTTCAT |
| *mPgk1* | TCACGGTGTTGCCAAAATGTC | TTGAAGTCCACCCTCATCACG |
| *mCyclophilin* | GGCAAATGCTGGACCAAACAC | TTCCTGGACCCAAAACGCTC |
| *hFABP4* | ACTGGGCCAGGAATTTGACG | AACTCTCGTGGAAGTGACGC |
| *hFABP5* | GTTCAGCAGCTGGAAGGAAGA | CATTGCGCCCATTTTTCGCA |
| *hPPARγ* | TATTCTCAGTGGAGACCGCC | AGGGCTTGTAGCAGGTTGTC |
| *h18s* | TTCCGACCATAAACGATGCCGA | GACTTTGGTTTCCCGGAAGCTG |

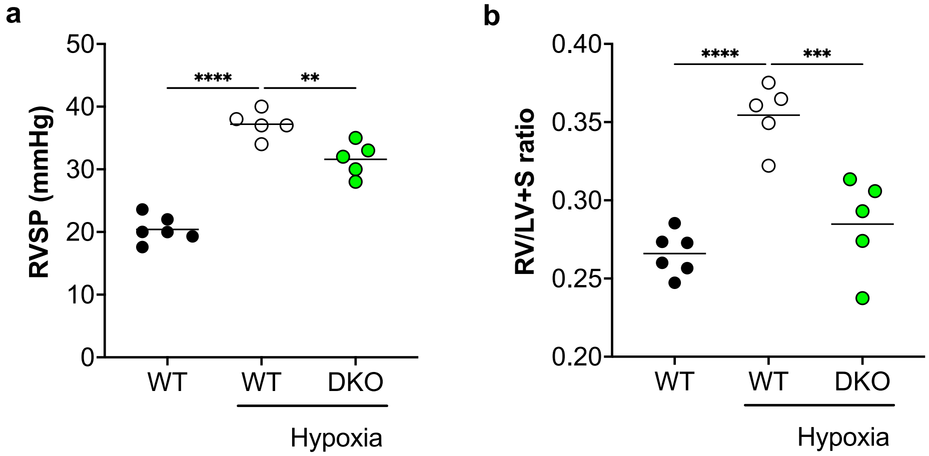


**Extended Data Fig 1. Fabp4-5$^{-/-}$ mice were protected from hypoxia-induced PH.**

(A) RVSP was reduced in Fabp4-5$^{-/-}$ mice compared to WT controls in hypoxia challenged condition. (B) RV hypertrophy was inhibited in Fabp4-5$^{-/-}$ mice compared to WT controls under hypoxia condition. ANOVA followed by Turkey post hoc analysis was used for statistical analysis (A, B). **P<0.01.

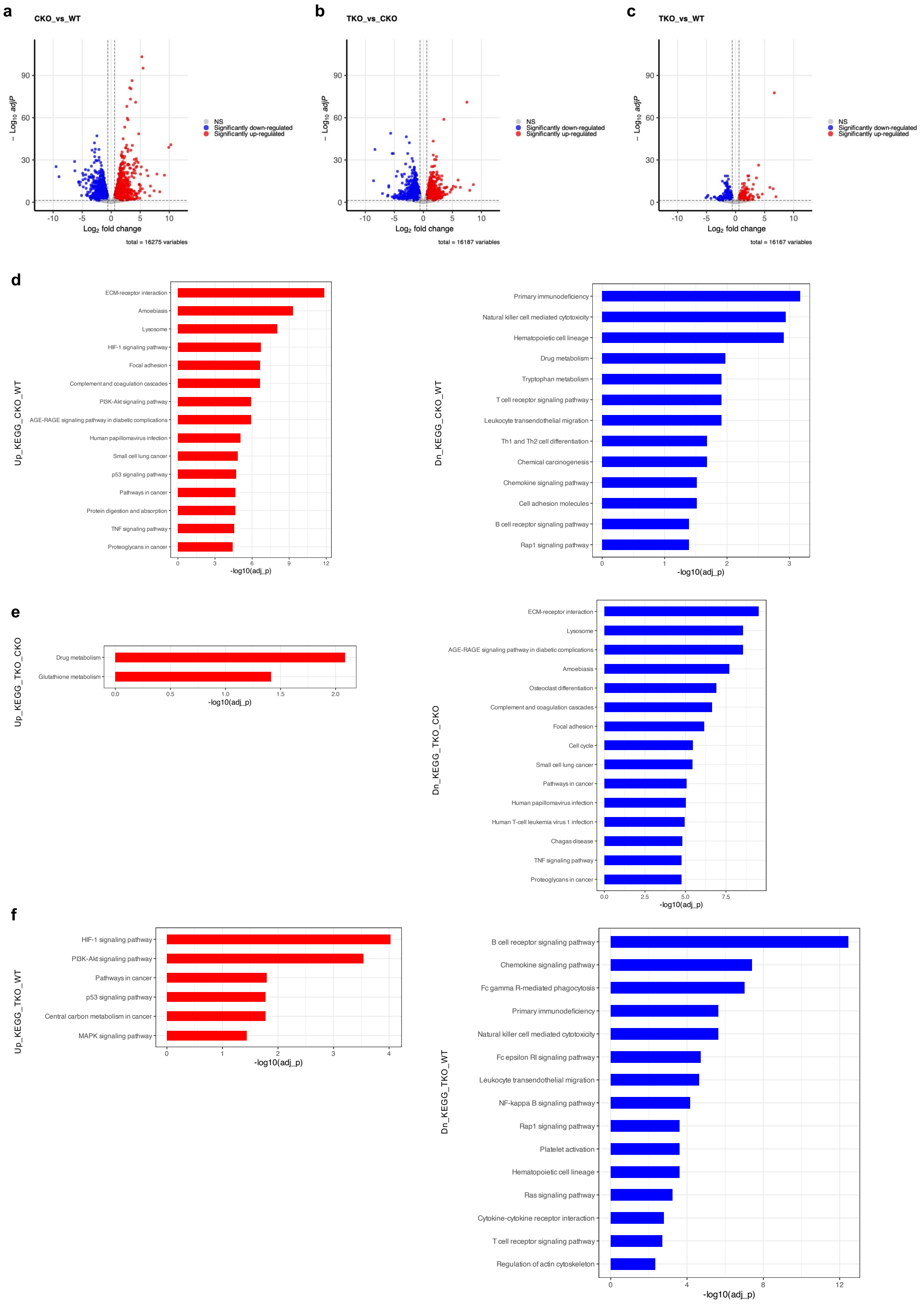
a
CKO_vs_WT
− Log10 adjP
Log2 fold change
NS
Significantly down-regulated
Significantly up-regulated
total = 16275 variables
b
TKO_vs_CKO
total = 16187 variables
c
TKO_vs_WT
total = 16167 variables
d
Up_KEGG_CKO_WT
ECM-receptor interaction
Amoebiasis
Lysosome
HIF-1 signaling pathway
Focal adhesion
Complement and coagulation cascades
PI3K-Akt signaling pathway
AGE-RAGE signaling pathway in diabetic complications
Human papillomavirus infection
Small cell lung cancer
p53 signaling pathway
Pathways in cancer
Protein digestion and absorption
TNF signaling pathway
Proteoglycans in cancer
-log10(adj_p)
Dn_KEGG_CKO_WT
Primary immunodeficiency
Natural killer cell mediated cytotoxicity
Hematopoietic cell lineage
Drug metabolism
Tryptophan metabolism
T cell receptor signaling pathway
Leukocyte transendothelial migration
Th1 and Th2 cell differentiation
Chemical carcinogenesis
Chemokine signaling pathway
Cell adhesion molecules
B cell receptor signaling pathway
Rap1 signaling pathway
e
Up_KEGG_TKO_CKO
Drug metabolism
Glutathione metabolism
Dn_KEGG_TKO_CKO
ECM-receptor interaction
Lysosome
AGE-RAGE signaling pathway in diabetic complications
Amoebiasis
Osteoclast differentiation
Complement and coagulation cascades
Focal adhesion
Cell cycle
Small cell lung cancer
Pathways in cancer
Human papillomavirus infection
Human T-cell leukemia virus 1 infection
Chagas disease
TNF signaling pathway
Proteoglycans in cancer
f
Up_KEGG_TKO_WT
HIF-1 signaling pathway
PI3K-Akt signaling pathway
Pathways in cancer
p53 signaling pathway
Central carbon metabolism in cancer
MAPK signaling pathway
Dn_KEGG_TKO_WT
B cell receptor signaling pathway
Chemokine signaling pathway
Fc gamma R-mediated phagocytosis
Primary immunodeficiency
Natural killer cell mediated cytotoxicity
Fc epsilon RI signaling pathway
Leukocyte transendothelial migration
NF-kappa B signaling pathway
Rap1 signaling pathway
Platelet activation
Hematopoietic cell lineage
Ras signaling pathway
Cytokine-cytokine receptor interaction
T cell receptor signaling pathway
Regulation of actin cytoskeleton

**Extended Data Fig 2. Transcriptomics analysis on the lungs of WT, CKO and TKO mice.**
The volcanos plots showing the gene expression change between WT and CKO mice (**A**), TKO and CKO mice (**B**), TKO and WT mice (**C**). The KEGG pathway analysis on the upregulated and downregulated DEGs between CKO and WT (**D**), TKO and CKO (**E**), as well as TKO and WT (**F**).

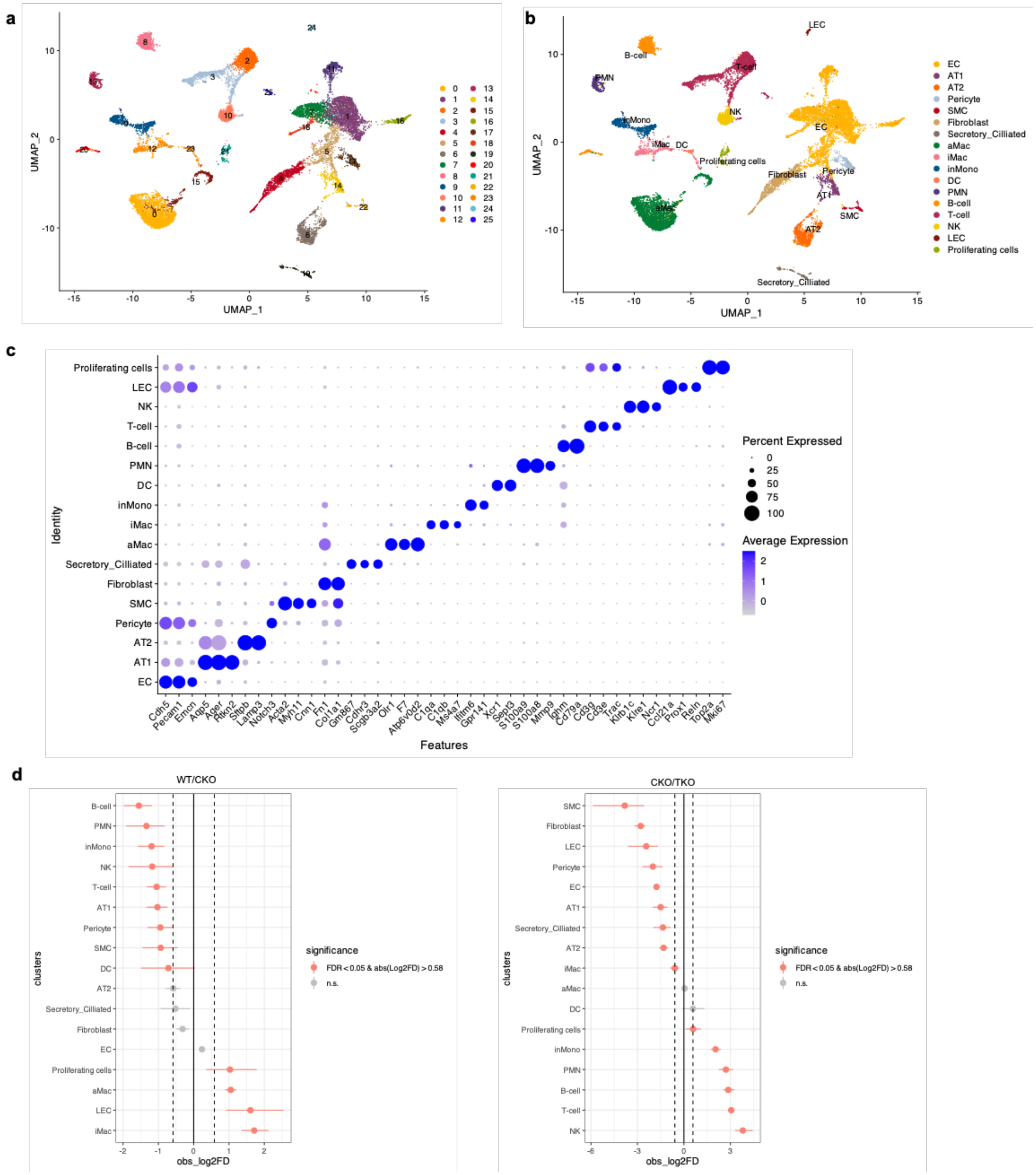


**Extended Data Fig 3**. **ScRNA-seq analysis on the lung cell populations from WT, CKO and TKO mice. (A)** A UMAP showing the integrated dataset. (**B**) A UMAP showing the individual group of cells for comparison. (C) A DotPlot showing the markers for each cell type. (D) Statistical analysis of the cell proportion change between WT and CKO on the left panel, and between CKO and TKO on the right panel.

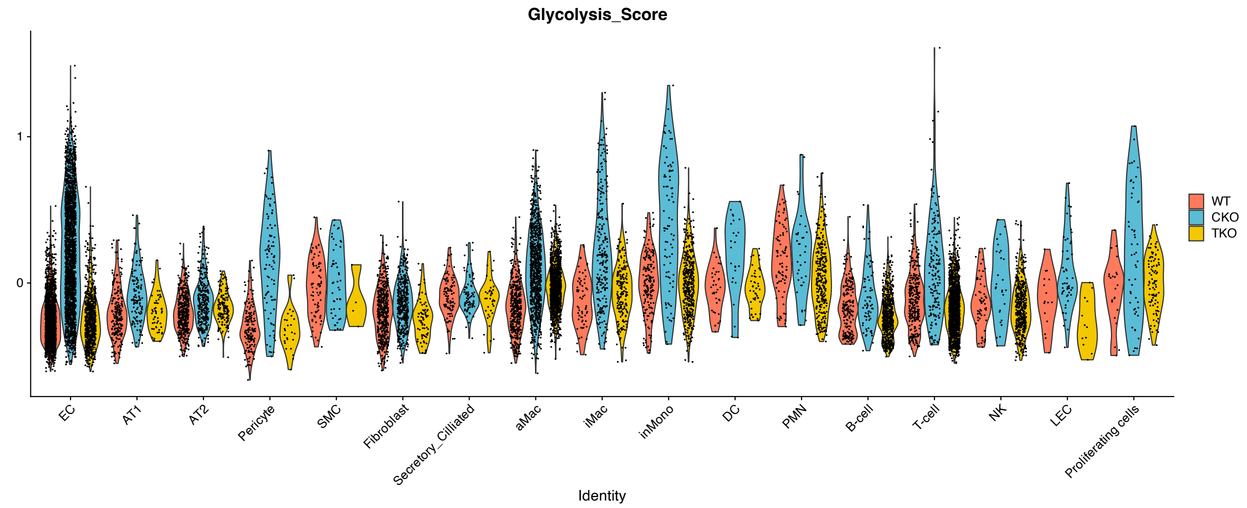


**Extended Data Fig 4. Glycolytic score calculation on the lung cell population amongst different mice strain.**

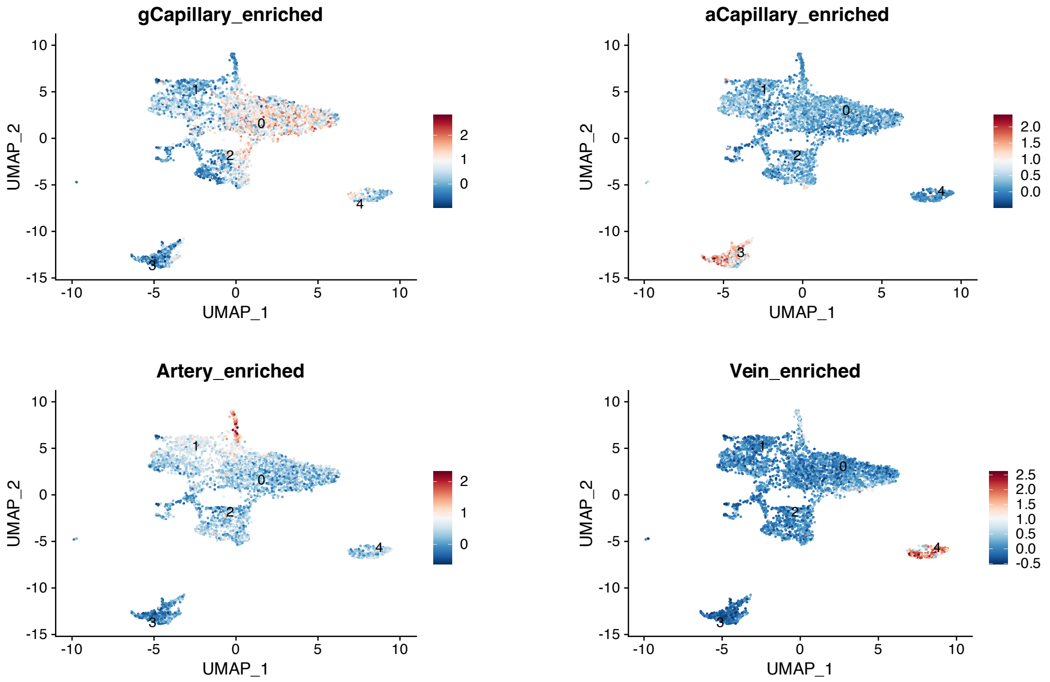


**Extended Data Fig 5. scRNA-seq analysis on the extracted EC subpopulation. (A)** Lung EC subpopulations markers enrichment analysis.

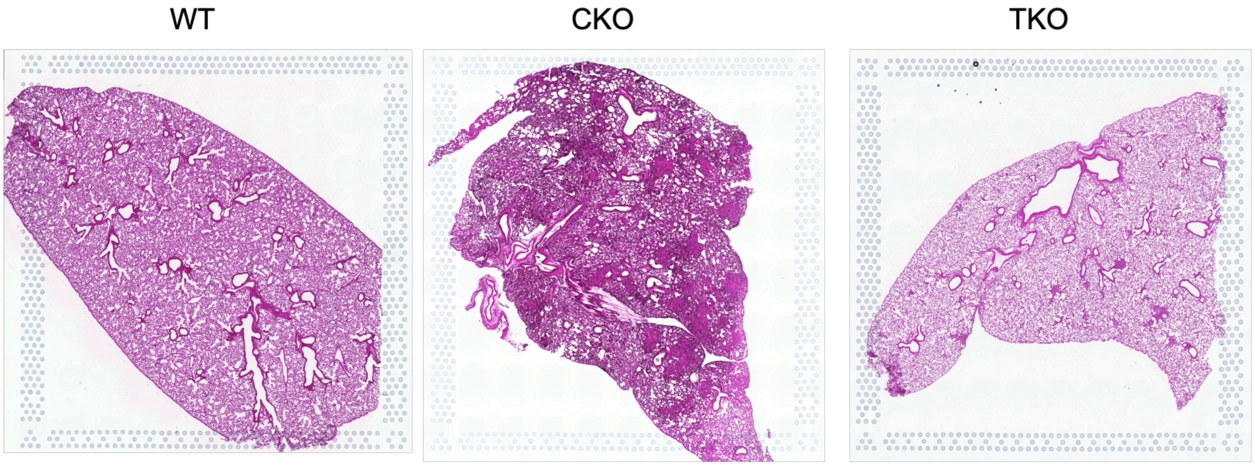


**Extended Data Fig 6. H&E staining for the tissue on Visium spatial transcriptomics analysis.** The images of WT and CKO were previously published. WT, CKO and TKO lungs slides were performed at the same time.

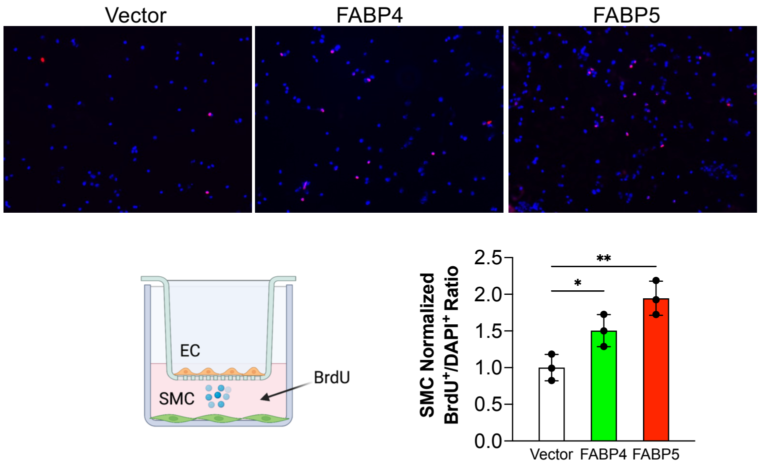


**Extended Data Fig 7. FABP4-5 promoted PASMCs proliferation.** Overexpression of FABP4 and 5 in hPAECs promoted PASMCs proliferation. ANOVA followed by Turkey post hoc analysis was used for statistical analysis. *P<0.05; **P<0.01.

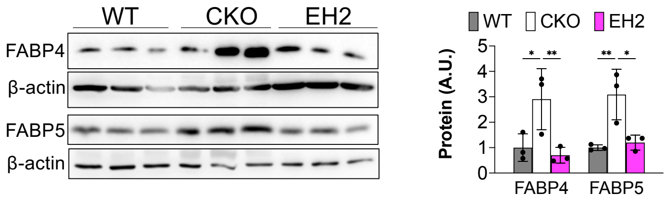


**Extended Data Fig 8. FABP4-5 upregulation in CKO mice is depended on HIF-2**α. ANOVA followed by Turkey post hoc analysis was used for statistical analysis. *P<0.05; **P<0.01.

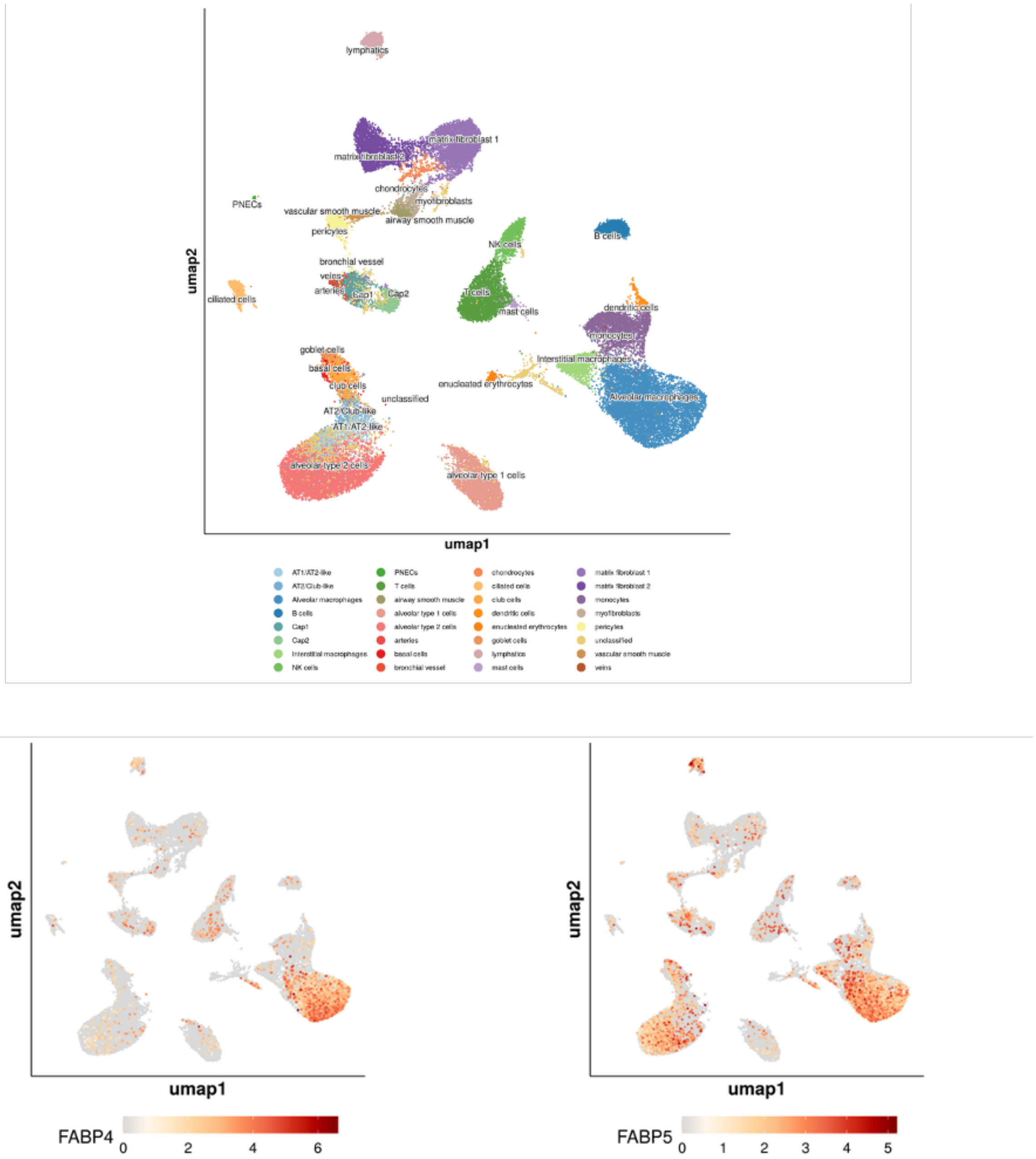


**Extended Data Fig 9. The expression pattern of FABP4 and 5 in healthy human lungs.** The scRNA-seq data from human lungs showed that FABP4 highly expressed in alveolar macrophage, whereas FABP5 strongly expressed in alveolar macrophage and AT2 cells in the healthy human lungs.

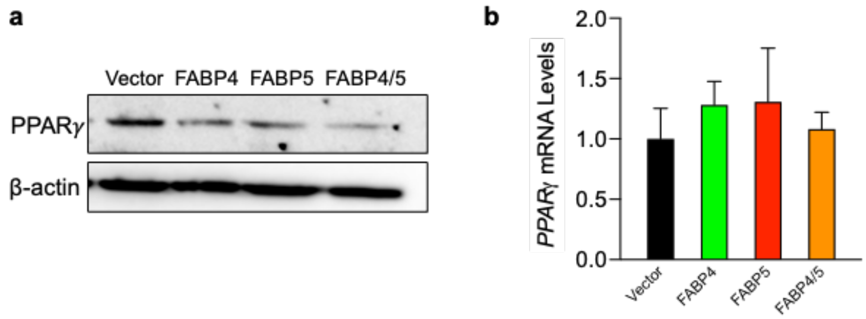


**Extended Data Fig 10. Overexpression of FABP4 and 5 reduce PPARγ protein but not mRNA expression in hPAECs.** (**A**) The western blotting analysis showed that PPARγ protein was reduced by FABP4 or FABP5 or both overexpression in hPAECs. (**B**) FABP4-5 did not affect PPARγ transcription via qPCR analysis. F4=FABP4, F5= FABP5, F4-5=FABP4 and FAPB5 combination.

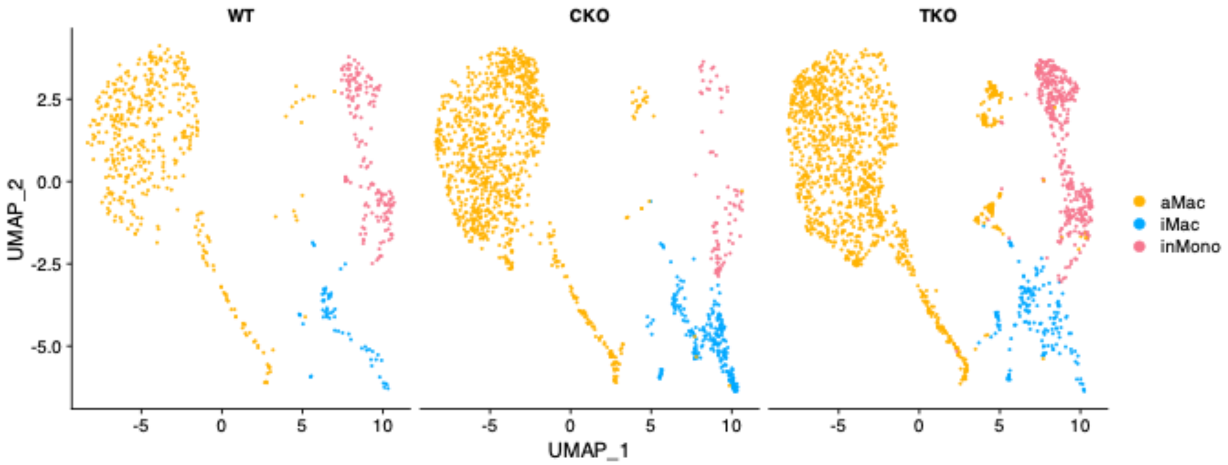


**Extended Data Fig 11. TKO mice does not affect macrophages population changes compared to CKO mice.** scRNA-seq data for the macrophage populations was extracted for re-clustering.
